# Strategies for the site-specific decoration of DNA origami nanostructures with functionally intact proteins

**DOI:** 10.1101/2021.07.01.450695

**Authors:** Joschka Hellmeier, René Platzer, Vanessa Mühlgrabner, Magdalena C. Schneider, Elke Kurz, Gerhard J. Schütz, Johannes B. Huppa, Eva Sevcsik

**Affiliations:** Institute of Applied Physics, TU Wien, Vienna, Austria; Center for Pathophysiology, Infectiology and Immunology, Institute for Hygiene and Applied Immunology, Medical University of Vienna, Vienna, Austria; Kennedy Institute of Rheumatology, University of Oxford, Oxford, UK

**Keywords:** DNA origami, DNA nanostructures, protein conjugation, functionalization, single molecule fluorescence microscopy, T-cell activation

## Abstract

DNA origami structures provide flexible scaffolds for the organization of single biomolecules with nanometer precision. While they find increasing use for a variety of biological applications, the functionalization with proteins at defined stoichiometry, high yield, and under preservation of protein function remains challenging. In this study, we applied single molecule fluorescence microscopy in combination with a cell biological functional assay to systematically evaluate different strategies for the site-specific decoration of DNA origami structures, focusing on efficiency, stoichiometry and protein functionality. Using an activating ligand of the T-cell receptor (TCR) as protein of interest, we found that two commonly used methodologies underperformed with regard to stoichiometry and protein functionality. While strategies employing tetravalent wildtype streptavidin for coupling of a biotinylated TCR-ligand yielded mixed populations of DNA origami structures featuring up to 3 proteins, the use of divalent (dSAv) or DNA-conjugated monovalent streptavidin (mSAv) allowed for site-specific attachment of a single biotinylated TCR-ligand. The most straightforward decoration strategy, via covalent DNA conjugation, resulted in a 3-fold decrease in ligand potency, likely due to charge-mediated impairment of protein function. Replacing DNA with charge-neutral PNA in a ligand conjugate emerged as the coupling strategy with the best overall performance in our study, as it produced the highest yield with no multivalent DNA origami structures and fully retained protein functionality. With our study we aim to provide guidelines for the stoichiometrically defined, site-specific functionalization of DNA origami structures with proteins of choice serving a wide range of biological applications.

---

DNA origami nanotechnology has emerged as a versatile tool for interrogating biological processes at a molecular and mechanistic level. By using short oligonucleotides (“staples”) to guide the folding of a long single stranded DNA scaffold, it is not only possible to program the shape of a DNA origami structure but also to arrange functional elements with nanometer resolution and precision^1^. Protein-decorated DNA origami structures have thus far been interfaced with cells as soluble agents^2,3^, attached to a solid substrate^4,5^ or anchored to a fluid supported lipid bilayer (SLB)^6,7^; biological questions addressed range from studying cellular signaling and adhesion processes^2–6,8,9^, to creating synthetic multienzyme cascades^10,11^ to more and more elaborate robotic DNA machines^12^.

The self-assembly of the DNA origami structures is typically straightforward. However, the challenge lies in their functionalization with proteins at defined stoichiometries with high yield while preserving protein function. A commonly applied method involves covalent conjugation of an oligonucleotide to a specific site within a protein of interest (reviewed in^13^) followed by hybridization to a complementary elongated staple strand (handle) on the DNA origami structure. The highly negatively charged DNA phosphate backbone, however, has been observed to compromise functionalization yield^14^ as well as protein functionality^15,16^. Genetically encoded protein tags^17,18^ or DNA-binding proteins^19^ allow for a defined stoichiometry and highly specific binding but require the generation of fusion proteins and often suffer from low coupling yields^15,20^, attributable to electrostatic repulsion at the coupling interface^14^. Alternatively, streptavidin (SAv) has been frequently used as a connector to attach biotinylated proteins to the DNA origami structure via a biotinylated handle^5,20–23^. This strategy has the advantage of shielding the protein from the negatively charged DNA. However, given the tetravalency of SAv for biotin-binding, single sites on the DNA origami may get functionalized with up to three proteins resulting in a stoichiometrically ill-defined product. While this may be acceptable for some applications, many mechanistic studies, e.g. those focusing on receptor-ligand interactions^2,3,6^, depend on the functionalization with not more than one protein at a specific site. We have recently circumvented this potential shortcoming of streptavidin by using divalent SAv (dSAv)^24^ as a connector to strictly avoid double or triple occupancies at a single modification site^6^.

As protein-functionalized DNA origami structures are becoming widely accessible research tools for the mechanistic study of diverse biophysical and cell biological processes, there is an increased need for robust methods to reliably produce high-quality DNA origami constructs. To date, however, systematic studies which provide the basis for the informed choice of a functionalization method are missing. With this work we offer a guideline for the site-specific, stoichiometrically defined functionalization of DNA origami structures with a protein of interest, with a particular focus on preserving full protein functionality. To this end, we systematically examined and optimized two commonly used approaches for the site-specific attachment of proteins to DNA origami structures, i.e. via commercially available streptavidin and DNA-protein conjugates, and introduced adaptations to these strategies to improve their performance. Specifically, we determined i) the yield of DNA origami structures functionalized with one (or more) proteins (functionalization efficiency), as well as ii) the number of proteins per functionalized DNA origami structure (functionalization stoichiometry) by using single molecule fluorescence microscopy. We chose an activating ligand of the T-cell receptor (TCR) for functionalization, which allowed us to assess iii) protein integrity by monitoring its stimulatory capacity in the presence of T-cells. In total, we compared seven different attachment strategies based on mono-, di- and tetravalent streptavidin as well as covalent conjugation via DNA and peptide nucleic acid (PNA) oligonucleotides (**Fig.1A**). While functionalization efficiencies for all tested strategies ranged from 67 to 74%, the use of tetravalent streptavidin did not yield stoichiometrically defined functionalization, which markedly affected the functional response to the ligand-decorated DNA origami constructs. Interestingly, the most straightforward approach for ligand coupling, using a DNA-ligand conjugate, markedly compromised ligand functionality. A strategy employing PNA instead of DNA for ligand conjugation gave rise to the best overall performance in our study, as it produced the highest yield of 74% with no multivalent DNA origami structures and fully retained protein functionality.

## Results and discussion

For the quantitative comparison of different functionalization strategies, we employed a recently introduced platform^6^ based on rectangular DNA origami tiles anchored to a fluid-phase supported lipid bilayer (SLB) via cholesterol-modified oligonucleotides^25^. DNA origami constructs were assembled from a 65×54 nm DNA origami tile^1^ featuring a centrally located and elongated staple strand to create a target for quantitative functionalization with proteins **(SI Fig. S1)**. As protein we selected for this study a monovalent single-chain antibody fragment of the variable domain derived from the TCRβ-reactive monoclonal antibody H57-597 (H57-scF_V_)^26^, which was shown to induce T-cell activation when displayed on SLBs^6,27^. We further equipped the H57-scF_V_ C-terminally with either a biotin ligase recognition sequence or an unpaired cysteine for the site-specific attachment to DNA origami structures, that would not interfere with TCR binding (**Fig. 1B**). All functionalization steps were carried out in solution followed by attachment of the fully assembled DNA origami constructs to a fluid SLB via cholesterol-modified oligonucleotides. Of note, all SLB-anchored DNA origami constructs exhibited free Brownian motion with a diffusion constant of ∼ 0.38 µm^2^/s (**SI Fig. S2, SI Table S1)**.

**Figure 1.**
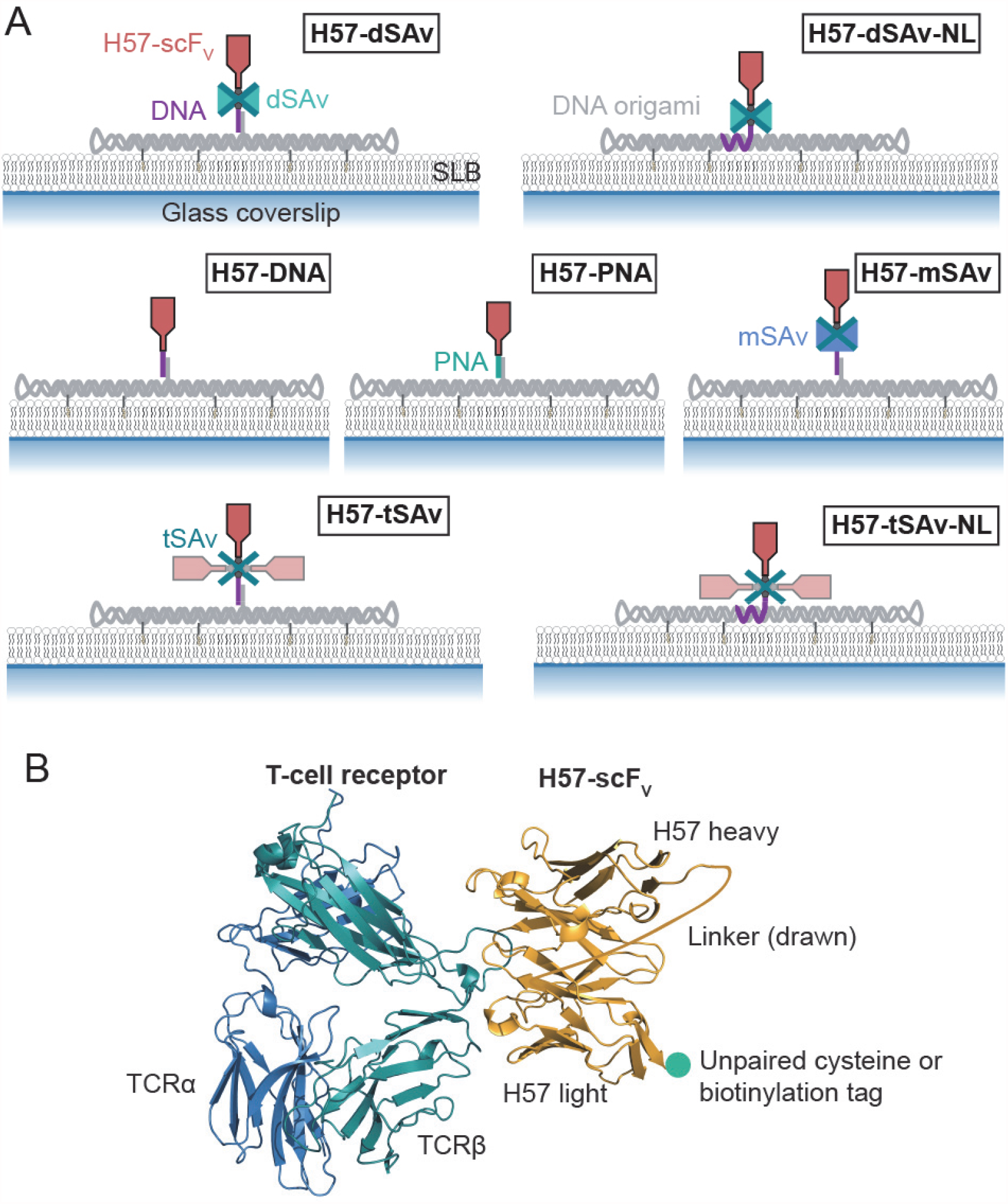
Strategies for site-specific functionalization of DNA origami with a single chain antibody fragment. **(A)** The single chain antibody fragment derived from the TCRβ-reactive mAb H57 (H57-scF_V_) was site-specifically attached to the DNA origami tile via different strategies. **(B)** Model based on a TCR–H57 Fab structure (PDB: 1NFD). At the C-terminus, the H57-scF_V_ was equipped with either an unpaired cysteine or an Avi-tag for site-specific biotinylation via birA.

To determine the efficiency of individual functionalization steps, we used single molecule two-color colocalization total internal reflection fluorescence (TIRF) microscopy **(Fig. 2A, SI Table 2)**. By varying molar ratios and incubation times, optimum conditions for each functionalization step were determined and then applied as the basis for subsequent steps **(SI Figs. S3-9; SI Tables S3-9)**. First, we determined the incorporation efficiency of the handle into the DNA origami tile. For this purpose, we employed a handle conjugated to Abberior Star 635P (DNA-AS635P) and labeled the DNA origami structure randomly with the DNA-intercalating fluorophore YOYO™-1 iodide (YOYO). Two color colocalization analysis of single molecule localizations recorded in the blue color channel (YOYO-1) with single molecule localizations recorded in the red color channel (DNA-AS635P) revealed an incorporation efficiency of ∼ 84% **(Fig. 2B)**, which is in good agreement with previously reported values for the center of a 2D DNA origami tile^28^. Note that this value represents a conservative estimate, as sub-stoichiometric labeling of the DNA-AS635P, fluorophore bleaching during recording and fluorophores present in the dark state were not taken into account.

**Figure 2.**
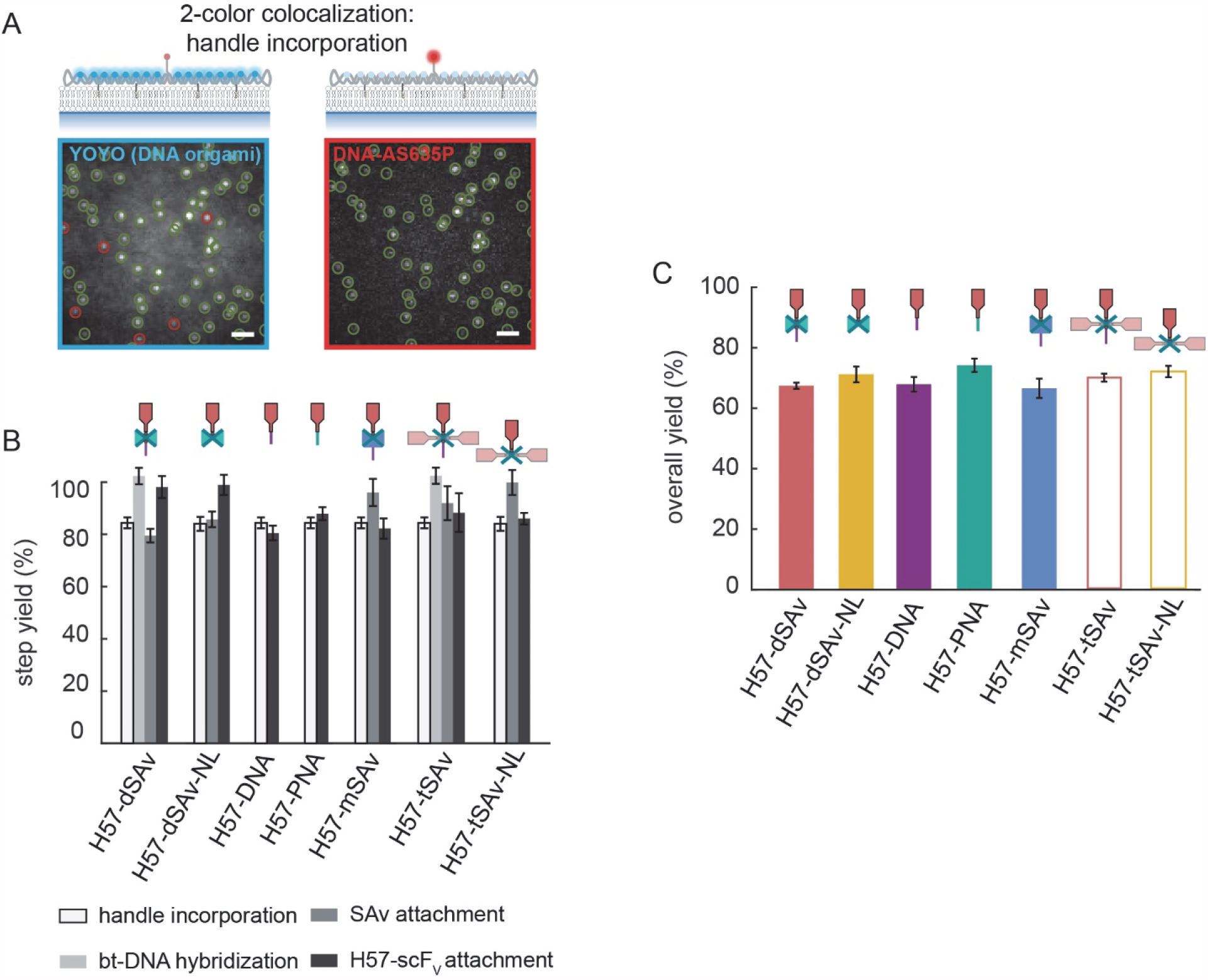
DNA origami functionalization efficiencies for the different site-specific attachment strategies. **(A)** Single molecule two-color colocalization TIRF imaging of DNA origami structures on a SLB was applied to determine the efficiency of each modification step. Determination of the incorporation efficiency of a fluorescently labeled handle (DNA-AS635P) in the DNA origami tile (labeled with the DNA-intercalating fluorophore YOYO) is shown as an example. Green open circles indicate signals detected in both color channels; red open circles indicate signals detected only in one channel. The percentage of colocalized signals in the blue (YOYO) and the red (DNA-AS635P) color channel amount to efficiency of handle incorporation. **(B)** The functionalization efficiency of each step was determined via two-color colocalization microscopy (see SI Figs. S3-9). **(C)** Functionalization yields of DNA origami structures with H57-scF_V_ are displayed. For each construct, data represent the mean of at least two independent experiments (± s.e.m.).

We have previously used a strategy where a biotinylated oligonucleotide was hybridized to the handle on the DNA origami tile followed by the attachment of divalent streptavidin (dSAv)^24^ and biotinylated Alexa Fluor 555 (AF555)-labeled H57-scF_V_ (**H57-dSAv, Fig. 1A**). This approach requires three additional functionalization steps ensuing the incorporation of the handle (**SI Fig. S3**); we determined the yield for each of these steps by two-color colocalization microscopy (**Fig. 2B**). By optimizing the molar ratios and incubation times (**SI Table S3**), we could increase the previously reported overall yield of ∼ 60%^6^ to ∼ 67% (**Fig. 2C, SI Table S10**). Furthermore, our results reveal that the availability of the handle and the efficiency of the subsequent modification steps contribute to a similar extent to the overall degree of functionalization. Note that two-color colocalization microscopy yields the efficiency of functionalization with at least one ligand. To assess the extent to which this equaled to *exactly one* ligand (strict 1:1 stoichiometry), we next determined the number of ligands per DNA origami construct by comparing the signal brightness of the construct to the brightness of a single AF555-labeled H57-scF_V_. As shown in **Fig. 3A**, virtually all localizations corresponded to single H57-scF_V_ molecules (**SI Fig. S10; SI Table S11**).

**Figure 3.**
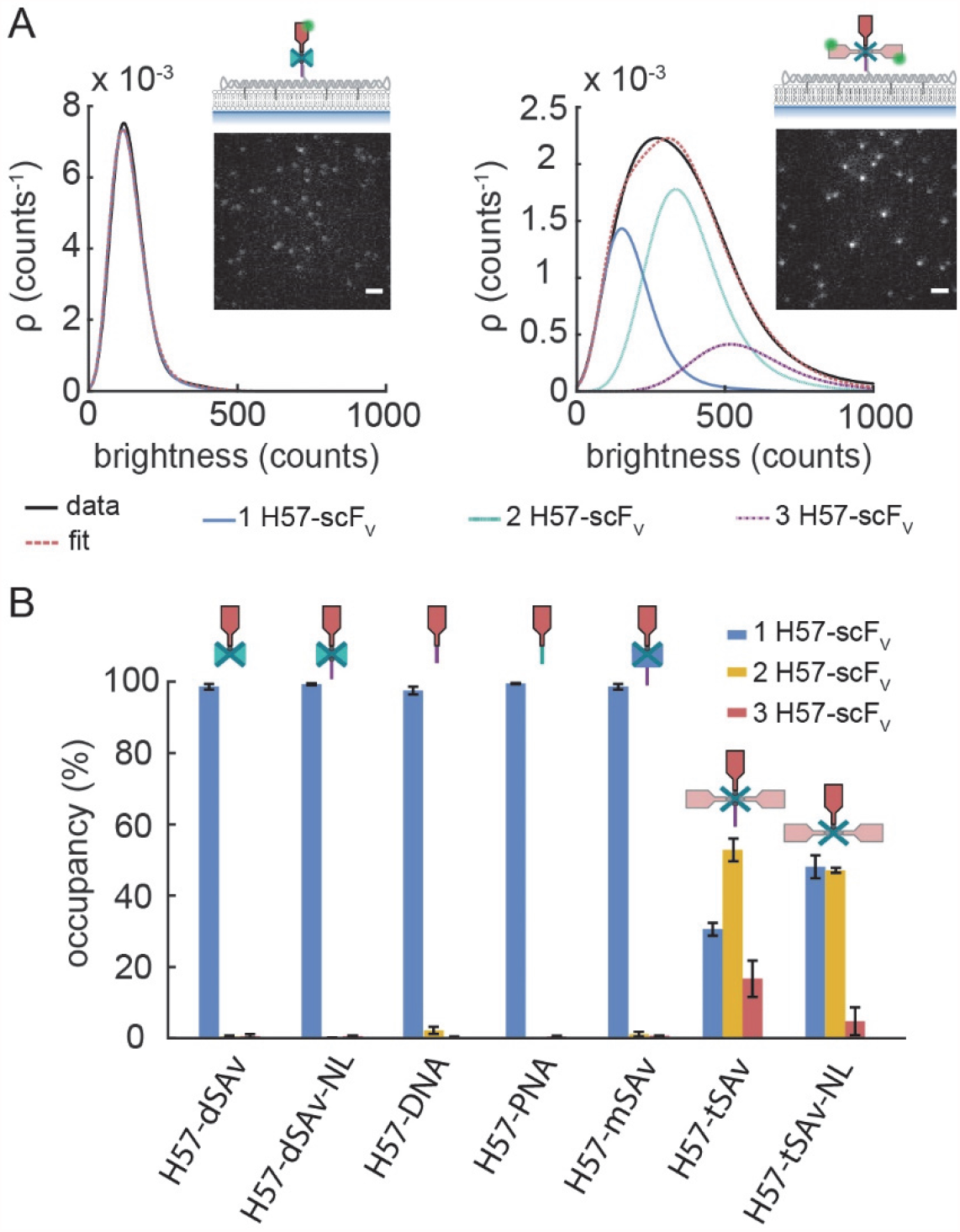
DNA origami functionalization stoichiometry for the different attachment strategies. **(A)** The number of H57-scF_V_s per DNA origami structure was determined via single molecule brightness analysis. Representative TIRF images (insets) and the corresponding brightness distributions ρ of biotinylated and AF555-conjugated scF_V_s bound via dSAv (H57-dSAv, left) and tSAv (H57-tSAv, right) to DNA origami structures on SLBs are shown. The detected signals were fitted and the brightness distribution was deconvolved into monomer and multimer contributions^29^ (see Methods section). Scale bar, 2 µm. **(B)** The percentage of detected (i.e. ligand-functionalized) DNA origami structures carrying 1, 2 or 3 H57-scF_V_s as determined from single-molecule brightness analysis. Means of at least two independent experiments are shown (± s.e.m.).

The functionalization strategy via dSAv as described above has several advantages. It reduces unwanted interactions between ligand and DNA, ensures a 1:1 stoichiometry of functionalization, and is cost-efficient and versatile, as the same biotinylated oligonucleotide can be used for introducing a functionalization site at various positions on the DNA origami tile by merely changing the position of the handle. The latter aspect, however, comes at a cost: stable hybridization of the biotinylated oligo to the handle requires at least 16 base pairs^30,31^ (we have used 17 nucleotides in this study), creating a double stranded DNA linker between DNA origami and dSAv with a length of ∼ 7 nm (**SI Fig. S11**). This, in turn, has several consequences that need to be considered: i) The DNA linker is connected to the DNA origami tile via 4 unpaired bases, thus conferring a certain degree of flexibility to the ligand which at the same time reduces the positional accuracy with respect to the DNA origami tile. ii) The linker increases the axial distance between the SLB-anchored DNA origami and the ligand, which may affect interactions that are sensitive to force and inter-membrane distance. In the construct **H57-dSAv-NL** (<u>n</u>o linker), we thus employed a short, biotinylated staple strand to directly attach dSAv to the DNA origami tile, which is expected to position the biotinylated ligand at a distance of ∼ 4 nm from the DNA origami surface, and thus permits a lower degree of motional freedom compared to the longer dsDNA linker present in construct H57-dSAv (**SI Fig. S4**). This attachment strategy resulted in a slightly higher functionalization yield (**Fig. 2B,C**).

Next, we determined whether the presence of the linker had an effect on the potency of the DNA origami constructs to activate T-cells. For this purpose, T-cells were confronted with SLBs presenting DNA origami constructs at different ligand surface densities together with His-tagged adhesion (ICAM-1) and co-stimulatory (B7-1) molecules as described previously^6^. For each experiment, the surface density of H57-scF_V_ on SLBs was assessed by relating the average fluorescence signal per area to the brightness of a single AF555-labeled H57-scF_V_ molecule. To monitor T-cell activation, T-cells were labeled with the calcium-sensitive dye Fura-2 AM, seeded onto DNA origami/SLB surfaces (**Fig. 4A**) and the level of intracellular calcium was assessed via ratiometric imaging (**SI Fig. S12**). The percentage of activated cells was determined for each SLB, plotted as dose-response curve (**Fig. 4B**) and fitted with Eq. 19 to determine H57-scF_V_densities at half-maximal response hereafter referred to as “activation threshold”. All fit parameters are listed in **SI Table S12**. For **H57-dSAv-NL** we determined an activation threshold of ∼ 3 H57-scF_V_per µm^2^ (**Fig. 4C**), similar to the value we had previously reported for **H57-dSAv**^6^, indicating that the dsDNA linker in the DNA origami construct did not markedly affect the potency of the TCR-ligand to activate T-cells.

**Figure 4:**
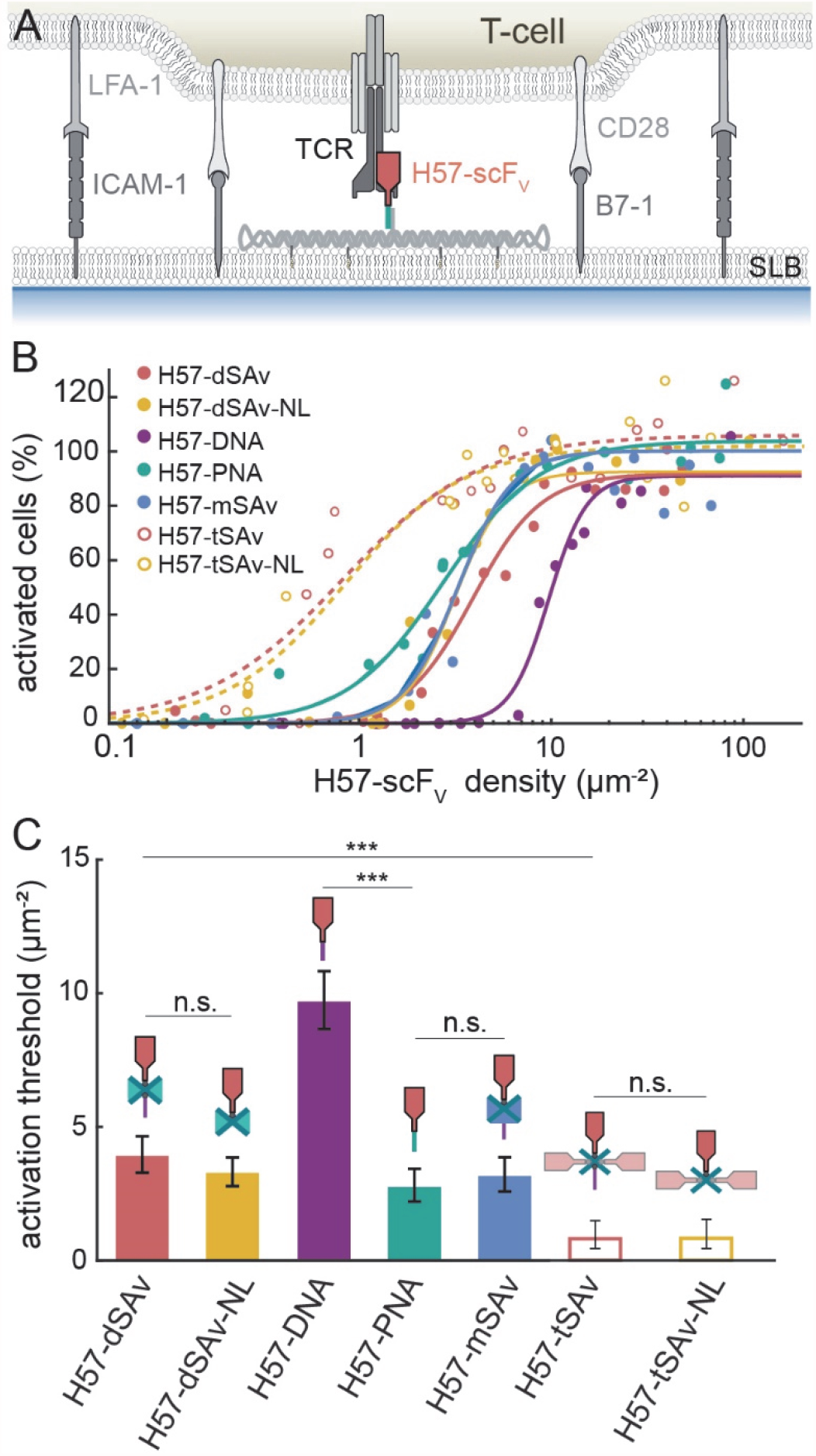
Ligand coupling strategy affects T-cell activation. **(A)** T-cells were interfaced with SLBs featuring ligand-decorated DNA origami constructs, adhesion (ICAM-1) and co-stimulatory molecules (B7-1). **(B)** Dose-response curves for T-cell activation mediated by H57-scF_V_ presented in the context of the different DNA origami constructs. Each data point corresponds to the percentage of activated cells determined in an individual experiment at a specific H57-scF_V_ density. Data were normalized to activation levels recorded for a positive control (=100%) involving the use of SLBs featuring His-tagged pMHC (His-pMHC) at a density of 150 µm^-2^ and His-tagged ICAM-1 and B7-1 at 100 µm^-2^. Dose-response curves were fitted with Eq. 19 (see Methods section) to extract activation thresholds **(C)**. For each dose-response curve, data are from at least two independent experiments involving T-cells isolated from two different mice. For details we refer to SI Table S12. Error bars represent the 95% confidence interval.

Although the individual steps for functionalizing the dSAv constructs were rather efficient, each modification step may be a source of error. Another drawback of strategies employing dSAv is that they do not allow the attachment of different ligands to create hetero-functional DNA origami structures. To address this limitation, we generated an H57-scF_V_-DNA conjugate by coupling a 17 base oligonucleotide to the free cysteine at the C-terminus of the H57-scF_V_ (see Methods section for details) and attached it directly to the complementary handle on the DNA origami tile via in-solution hybridization (**H57-DNA, Fig. 1A, SI Fig. S5**) at an efficiency of ∼ 80%, yielding a total coupling efficiency of ∼ 68 % (**Fig. 2B,C**). We found, however, that ligand functionality was considerably reduced, as evidenced by a 3-fold increased activation threshold (10 µm^-2^ compared to 3 µm^-2^ for H57-dSAv-NL, **Fig. 3B,C**). Indeed, conjugation of DNA oligonucleotides to proteins has been reported to compromise hybridization efficiency^14^ as well as protein functionality^15,16^, possibly due to electrostatic interactions between positively charged patches on the protein and the negatively charged DNA. Considering the rather small size of H57-scF_V_ (27 kDa) it is plausible that direct coupling of the highly negatively charged DNA oligonucleotide causes local pH-changes which could interfere with TCR binding. Indeed, while soluble, unmodified H57-scF_V_ binds the TCR with high affinity at ∼ 95% labeling efficiency^26^, which amounts to a surface density of ∼95 H57-scF_V_-labled TCRs per µm^2^ on the T-cell surface, we found that TCR staining by H57-DNA was markedly reduced (∼ 20 H57-scF_V_-labled TCRs per µm^2^) (**SI Fig. S13**).

Considering the detrimental effect of the negatively charged DNA tag on ligand activity, we decided to use a PNA (peptide nucleic acid) oligo as a charge-neutral alternative ^32^. The PNA backbone is composed of repeating peptide-like amide units (N-(2-aminoethyl) glycine), thereby supporting high-affinity PNA-DNA duplex formation due to the absence of inter-strand electrostatic repulsion. In a recent study, to our knowledge the only one employing PNA for functionalization of DNA origami thus far, a ligand conjugated with a PNA oligonucleotide of only 9 bases was efficiently coupled to a DNA handle on a DNA origami structure ^3^. Indeed, H57-scF_V_functionality could be completely restored by substituting the DNA oligo with PNA (**H57-PNA, SI Fig. S6**), giving rise to an activation threshold of 2.5 H57-scF_V_ molecules per µm^2^ (**Fig. 4B,C**) as well as a higher hybridization efficiency of 88%, and thus a total functionalization efficiency of ∼ 74%. (**Fig. 2B,C; SI Table S10**). Accordingly, soluble H57-PNA labeled TCRs on the T-cell surface with a similar efficiency when compared to unconjugated or biotinylated H57-scF_V_ (**SI Fig. S13**).

To combine the advantages of streptavidin-based strategies (i.e. use of a biotinylated protein, avoidance of electrostatic interactions between DNA and the protein of interest) and hybridization-based strategies (i.e. creation of hetero-functional DNA origami), we introduced a DNA-conjugated monovalent tetrameric streptavidin (mSAv) as a spacer protein, which was attached to the DNA origami tile via hybridization to the handle, followed by coupling of a site-specifically biotinylated H57-scF_V_ to the mSAv **(H57-mSAv, SI Fig. S7)**. Indeed, using mSAv to shield the H57-scF_V_ from the negatively charged DNA fully restored ligand functionality (**Fig. 4B,C**) at a total coupling yield of 67% (**Fig. 2C**) and a defined 1:1 stoichiometry (**Fig. 3B**).

The attachment strategies described so far require biochemical expertise and instrumentation which may not be accessible in every lab. While ensuring stoichiometrically well-defined binding, neither mSAv nor dSAv are readily available. Instead, commercially available tetravalent streptavidin (tSAv) has often been used, and while it offers in principle three sites for ligand binding, only one may be accessible due to steric hindrance when surface-attached^33^. Differences in the cellular response to a ligand presented via mSAv and tSAv have been observed yet attributed to the presence of a flexible linker in the mSAv construct^7^. To assess biochemical and functional consequences of tSAv valency in more detail, we designed the construct **H57-tSAv** in analogy to H57-dSAv, but replaced the divalent with tetravalent streptavidin (**SI Fig. S8**). While showing similar functionalization efficiency (**Fig. 2B,C**), this strategy yielded a mixed population, with detected DNA origami constructs featuring one (∼ 30%), two (∼ 53%) and three ligands (∼ 17%) as determined via single molecule brightness analysis (**Fig. 3A,B, SI Table S11**).

In a previous study, we had found that the lateral spacing of H57-scF_V_ dramatically affected T-cell activation^6^. Presentation of two H57-scF_V_ ligands at spacings below 20 nm resulted in substantially increased potency compared to ligand spacings of ≥ 48 nm, as enforced by the DNA origami tiles also used in the present study (**Fig. 4B,C**). Indeed, the H57-tSAv construct yielded an activation threshold of ∼ 0.3 µm^-2^, almost one order of magnitude lower than the strictly monovalent constructs and similar to the value we had previously determined for divalent constructs. It hence appears likely that at least two of the three tSAv-bound H57-scF_V_s, that are maximally available per DNA origami, participate in TCR binding.

The binding geometry of H57-scF_V_ to the TCR^26^ renders the ligand perpendicular to the SLB, i.e. H57-scF_V_ bound to the trans binding pocket of tSAv, most effective for TCR binding. In an attempt to decrease the availability of the two biotin binding pockets adjacent to the one used for attachment to the DNA origami tile, we omitted the dsDNA linker in construct **H57-tSAv-NL** to attach tSAv closer to the DNA origami surface (**SI Fig. S9**). While this configuration significantly reduced triple occupancies (to ∼ 5%), still half of all signals originated from DNA origami tiles carrying two H57-scF_V_s (**Fig. 3B**). Accordingly, **H57-tSAv-NL** activated T-cells with high potency (**Fig. 4B**), indicating that our efforts to decrease the availability of the second tSAv-bound H57 for TCR binding were not successful. While functionalization with larger proteins would likely increase the fraction of monovalent DNA origami structures due to steric hindrance, we consider it unlikely that a strictly monovalent population can be achieved using tSAv for attachment. Given that exact placement of proteins within DNA origami structures is a primary reason for their use in the first place, this obvious lack of experimental control diminishes the usefulness of tetravalent streptavidin in many biological and biophysical applications. In fact, we have previously observed that the presence of only 0.5% DNA origami structures carrying two H57-scF_V_s at a distance of 10 nm instead of a single H57-scF_V_ dominates T-cell activation behavior^6^ with obvious consequences for the kind of conclusions that can be drawn from such experiments.

## Conclusion

We have here systematically evaluated and optimized strategies for site-specific protein functionalization of DNA origami structures, with the aim of presenting guidelines tailored towards different experimental capacities and requirements. Focusing particularly on functionalization stoichiometry and protein functionality, we found that two commonly used methodologies underperformed with regard to these critical aspects: covalent conjugation of a DNA oligonucleotide and subsequent hybridization to the DNA origami structure resulted in a 3-fold decreased ligand potency. For strategies employing tetravalent streptavidin, on the other hand, the majority of DNA origami structures was functionalized with more than one biotinylated protein at a single modification site, rendering this approach inadequate if the anticipated experiment mandates a defined stoichiometry of target protein to allow for a conclusive outcome.

Using charge-neutral PNA oligos for protein conjugation emerged as the strategy with the best overall performance in our study, as it produced the highest coupling yield of 74% with no multivalent DNA origami structures and fully retained protein functionality. Moreover, PNA-based functionalization is well suited for creating hetero-functionalized DNA origami that carry several different proteins of interest. A versatile and less cost-intensive alternative to PNA conjugation at a slightly lower yield was the use of DNA-conjugated mSAv and a biotinylated protein, where SAv acted as spacer to the negatively charged DNA. Similar to direct protein conjugation of DNA or PNA oligonucleotides, this strategy allows the generation of hetero-functional DNA origami structures by pre-incubating the proteins of interest with mSAv prior to hybridization. In conclusion, PNA- and mSAv-based strategies are highly valuable for the site-specific protein functionalization of DNA origami structures and for serving a wide range of biological applications.

## Materials & Methods

### Assembly of DNA origami tiles

DNA origami structures were assembled in a single folding reaction carried out in a test tube (AB0620, ThermoFisher Scientific) with 10 µl of folding mixture containing 10 nM M13mp18 scaffold DNA (New England Biolabs), 100 nM unmodified oligonucleotides (Integrated DNA technologies), either 100 nM biotin-modified oligonucleotides (Biomers) for direct hybridization to the DNA origami tile (H57-dSAv-NL, H57-tSAv-NL) or 500 nM biotinylated oligonucleotides (Biomers) for external hybridization (H57-dSAv, H57-tSAv) and folding buffer (5 mM Tris (AM9855G, ThermoFisher Scientific), 50 mM NaCl (AM9759, ThermoFisher Scientific), 1 mM EDTA (AM9260G, ThermoFisher Scientific), 12.5 mM MgCl2) (AM9530G, ThermoFisher Scientific)). Oligonucleotide sequences are shown in the **SI Appendix, Table S13-15**. At the site chosen for ligand attachment, a staple strand was elongated at its 3’-end with 21 bases (H57-DNA, H57-PNA, H57-mSAv, H57-dSAv, H57-tSAv). At sites chosen for cholesterol anchor attachment, staple strands were elongated at 5’-end with 25 bases, respectively. DNA origami were annealed using a thermal protocol (90°C, 15 min; 90°C – 4°C, 1°C min^-1^; 4°C, 6h) and purified using 100kDa Amicon ®Ultra centrifugal filters (UFC510096, Merck). DNA origami were stored up to 4 weeks at -20°C.

### Functionalization of DNA origami tiles

In the following, assembly strategies are given at optimal conditions for each construct. For further details regarding the individual functionalization steps, we refer to **SI Figs. S3-9**. Functionalized DNA origami constructs were not stored, but used for experiments at the same day.

### Construct 1: H57-dSAv

For H57-dSAv, DNA origami were functionalized in a three – step assembly process (SI Fig. S3). A biotinylated oligonucleotide was hybridized at 5x molar excess to a protruding elongated staple strand during the initial thermal annealing process of the DNA origami tile followed by purification using 100kDa Amicon®Ultra centrifugal filters (Merck). In a next step, DNA origami structures were incubated with a 10x molar excess of dSAv for 30 min at 24°C and excessive dSAv was removed using 100kDa Amicon®Ultra centrifugal filters. As a last step, AF555-conjugated and site-specifically biotinylated H57-scF_*V*_ was added at a 10x molar excess for 60 min at 24°C. Finally, H57-scF_*V*_ not bound to DNA origami structures was removed using 100kDa Amicon®Ultra centrifugal filters.

### Construct 2: H57-dSAv-NL

For H57-dSAv-NL, DNA origami were functionalized in a three-step assembly process (SI Fig. S4). At the site chosen for ligand attachment, the staple strand was modified with a biotin group and added at a 10x molar excess to the DNA origami folding mix during the initial thermal annealing process followed by purification. Next, DNA origami were incubated with a 10x molar excess of dSAv for 60 min at 24°C and excessive dSAv was removed using 100kDa Amicon®Ultra centrifugal filters. AS635P-conjugated and site-specifically biotinylated H57-scF_*V*_ was added at a 10x molar excess for 60 min at 24°C. Finally, excessive H57-scF_*V*_ was removed using 100kDa Amicon®Ultra centrifugal filters.

### Construct 3: H57-DNA

For H57-DNA, DNA origami were functionalized in a two-step assembly process (SI Fig. S5). Here, the AF555-conjugated H57-scF_*V*_ was site-specifically modified with a complementary DNA strand (DNA-H57) (see below) and hybridized to the elongated staple strand on the DNA origami tile. For functionalization, DNA origami were incubated with a 10x molar excess of DNA-H57 for 60 min at 35°C and cooled down to 4°C at 1°C min^-1^. Excessive DNA-H57 was removed during a final purification step using 100kDa Amicon®Ultra centrifugal filters.

### Construct 4: H57-PNA

For H57-PNA, DNA origami were functionalized in a two-step assembly process (SI Fig. S6). For this purpose, the AF555-conjugated H57-scF_*V*_ was site-specifically modified with a complementary PNA strand (PNA-H57) and hybridized to the elongated staple strand on the DNA origami tile. For functionalization, DNA origami were incubated with a 3x molar excess of PNA-H57 for 60 min at 35°C and cooled down to 4°C at 1°C min^-1^. Excessive PNA-H57 was removed during a final purification step using 100kDa Amicon®Ultra centrifugal filters.

### Construct 5: H57-mSAv

For H57-mSAv, DNA origami were functionalized in a three-step assembly process (SI Fig. S7). For this purpose, a complementary DNA oligo was conjugated to a free cysteine on mSAv (DNA-mSAv) (see below) and hybridized to the elongated staple strand on the DNA origami tile. For this, DNA origami were incubated with a 3x molar excess of DNA-mSAv for 60 min at 35°C and cooled down to 4°C at 1°C min^-1^. Excessive DNA-mSAv was removed using 100kDa Amicon®Ultra centrifugal filters. Finally, AF555-conjugated and site-specifically biotinylated H57-scF_*V*_ was added at a 10x molar excess for 60 min at 24°C, followed by a final purification step to remove excessive H57-scF_*V*_.

### Construct 6: H57-tSAv

For H57-tSAv, DNA origami were functionalized in a three-step assembly process (SI Fig. S8) similarly to H57-dSAv. A biotinylated oligonucleotide was hybridized at 5x molar excess to the complementary elongated staple strand on the DNA origami tile during the initial thermal annealing process followed by purification. In a next step, DNA origami were incubated with a 10x molar excess of tSAv for 30 min at 24°C. Excessive tSAv was removed using 100kDa Amicon®Ultra centrifugal filters. Further, AF555-conjugated and site-specifically biotinylated H57-scF_*V*_ was added at a 10x molar excess for 60 min at 24°C. Finally, excessive H57-scF_*V*_ was removed using 100kDa Amicon®Ultra centrifugal filters.

### Construct 7:H57-tSAv-NL

For H57-tSAv-NL, DNA origami were functionalized in a three-step assembly process (SI Fig. S9) in analogy to H57-dSAv-NL.

### Preparation of functionalized planar SLBs

For DNA origami characterization, vesicles containing 100% 1-palmitoyl-2-oleoyl-sn-glycero-3-phosphocholine (POPC) (Avanti Polar Lipids) were prepared at a total lipid concentration of 0.5mg ml^-1^ as described^26^ in 10x Dulbecco’s phosphate-buffered saline (PBS) (D1408-500ml, Sigma Aldrich).

Glass coverslips (# 1.5, 24×60 mm, Menzel) were plasma cleaned for 10 min and attached with the use of dental imprint silicon putty (Picodent twinsil 22, Picodent) to Lab-Tek™ 8-well chambers (ThermoFisher Scientific), from which the glass bottom had been removed^34^. Coverslips were incubated with a fivefold diluted vesicle solution for 10 min, before they were extensively rinsed with PBS (D1408-500ML, Sigma Aldrich). For functionalization, SLBs were first incubated for 60 min with cholesterol-modified oligonucleotides (Integrated DNA technologies) complementary to the elongated staple strands at the bottom side of the DNA origami and then rinsed with PBS. DNA origami were incubated on SLBs in PBS + 1% BSA (A9418-10G, Sigma Aldrich) for 60 min. For T-cell activation experiments, vesicles containing 98% 1-palmitoyl-2-oleoyl-sn-glycero-3-phosphocholine (POPC) and 2% 1,2-dioleoyl-sn-glycero-3-[N(5-amino-1-carboxypentyl) iminodiacetic acid]succinyl[nickel salt] (Ni-DOGS NTA) (Avanti Polar Lipids) were used and SLBs were formed as described above. Upon incubation of DNA origami on SLBs, His_10_-tag ICAM-1 (50440-M08H, Sino Biological) (270 ng ml^-1^) and His_10_-tag B7-1 (50446-M08H, Sino Biological) (130 ng ml^-1^) were incubated for 75 min at 24°C and then rinsed off with PBS. PBS was replaced with HBSS for single molecule imaging (H8264-500ML, Sigma Aldrich) and HBSS + 2% FBS for T-cell activation experiments.

### Total internal reflection fluorescence (TIRF) microscopy

TIRF microscopy experiments were performed on a home-built system based on a Zeiss Axiovert 200 microscope equipped with a 100x, NA=1.46 Plan-Apochromat objective (Zeiss). TIR illumination was achieved by shifting the excitation beam parallel to the optical axis with a mirror mounted on a motorized table. The setup was equipped with a 488 nm diode laser (iBeam smart 488, Toptica), a 532 nm diode-pumped solid state (DPSS) laser (Spectra physics Millennia 6s) and a 647 nm diode laser (Obis LX 647, Coherent). Laser lines were overlaid with an OBIS Galaxy beam combiner (Coherent).

Direct analog laser modulation (488 and 647 nm) or an Acousto-optic modulator (Isomet) (532 nm) were used to adjust laser intensities (1-3kW cm^-2^) and timings using an in-house developed package implemented in LABVIEW (National Instruments). A dichroic mirror (Di01-R405/488/532/635-25×36, Semrock) was used to separate excitation and emission light. Emitted signals were split into two color channels using an Optosplit II image splitter (Cairn) with a dichroic mirror (DD640-FDi01-25×36, Semrock) and emission filters for each color channel (FF01-550/88-25, ET 570/60, ET 675/50, Chroma) and imaged on the same back-illuminated EM-CCD camera (iXon Ultra, DU897, Andor).

### Determination of functionalization efficiencies via two-color colocalization TIRF microscopy

To determine the efficiency of a particular functionalization step, two fluorescently labeled interaction partners were used and the efficiency of functionalization was determined via two-color colocalization analysis. All fluorescently labeled interaction partners are listed in SI Table S2). After the particular functionalization, DNA origami constructs were anchored to SLBs as described above and positions of diffraction-limited spots were determined in both color channels. Single molecules were localized and corrected for chromatic aberrations as described^35^. Detected signal positions were counted as colocalized if signals were within a distance of 240 nm. The starting construct was always assigned color channel 1 (DYE 1), the functionalization to be attached at the particular step was assigned color channel 2 (DYE 2). The fraction of colocalized signals, *f*_*coloc XY*_, where × denotes the construct number and Y the functionalization step (SI Figs. S3-9), was determined by relating the number of signals in the second color channel (DYE 2) that colocalized with a signal in the first color channel (DYE 1), *N*_*coloc XY*_, to the number of detected signals in the first color channel, *N*_*total XY*_ (Eq. 1).

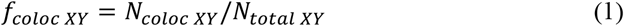

Based on Eq. 1, the functionalization efficiency of each functionalization step was determined for each construct separately.

Each functionalization step, in turn, was optimized with regard to molar ratios and incubation times via two-color colocalization experiments (SI Tables S3-9). Once a functionalization step was optimized, these conditions were used as the basis for subsequent steps.

### Handle incorporation

A staple strand fluorescently labeled at its 3’-end with AS635P was used in the assembly process, and DNA origami were pre-stained with YOYO™-1 iodide (YOYO) at a concentration of 1 µg ml^-1^ for 45 min at 24°C. Excessive YOYO was removed using 100 kDa Amicon®Ultra centrifugal filters and DNA origami-bearing SLBs were produced as described above. The fraction of incorporated elongated staple strands, *f*_*coloc* 11_, was determined by relating the number of signals in the red color channel (DNA-AS635P) that colocalized with a signal in the blue color channel (YOYO), *N*_*coloc* 11_, to the number of detected signals in the blue color channel, *N*_*total* 11_ (Eq. 2).

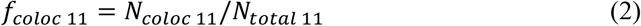

The incorporation of biotinylated staple strand was determined similarly, with the exception that a handle modified with an Alexa Fluor 647 at its 5’-end and a biotin at its 3’-end (AF647-DNA-bt) was used (Eq. 3).

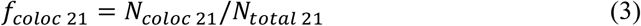

### Construct 1: H57-dSAv

The second modification step (following incorporation of the handle) was the hybridization of a biotinylated oligo to the handle. The fraction of DNA origami carrying a biotin modification, *f*_*coloc* 12_, was determined. For this, DNA origami were assembled with a handle modified with AS635P at the 3’-end and biotin at the 3’-end (bt-DNA-AS635P). DNA origami were pre-stained with YOYO as described above. By relating the number of signals in the red color channel (bt-DNA-AS635P) that colocalized with a signal in the blue color channel (YOYO), *N*_*coloc* 12_, to the number of detected blue signals, *N*_*total* 12_, *f*_*coloc* 12_ could be determined (Eq. 4).

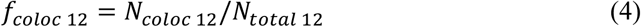

Next, the fraction of DNA origami carrying dSAv for further ligand attachment, *f*_*coloc* 13_, was determined. For this, DNA origami labeled with bt-DNA-AS635P were incubated with AF555-conjugated dSAv. The number of green signals (dSAv) that colocalized with a red signal (bt-DNA-AS635P), *N*_*coloc* 13_, was divided by the number of red signals, *N*_*total* 13_. After correction for the fraction of DNA origami carrying neither biotin nor dSAv, we arrive at the fraction of DNA origami carrying a dSAv (Eq. 5).

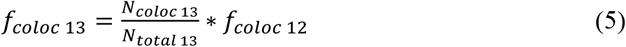

Finally, the fraction of DNA origami carrying a TCR ligand, *f*_*coloc* 14_, was determined by using DNA origami labeled with bt-DNA-AS635P and AF555-conjugated H57-scF_*V*_. By evaluating the number of green signals (H57-scF_V_) that colocalized with a red signal (bt-DNA-AS635P), *N*_*coloc* 14_, divided by the number of red signals (*N*_*total* 14_) and corrected for the fraction of unoccupied DNA origami, the fraction of DNA origami functionalized with H57-scF_V_ could be determined (Eq. 6).

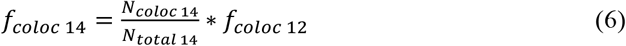

### Construct 2: H57-dSAv-NL

The second modification step (following incorporation of the biotinylated handle) concerned the attachment of AF555-labeled dSAv (dSAv-AF555). For this, AF647-DNA-bt was employed as handle. The fraction of biotin-bound dSAv was evaluated by dividing the number of green signals (dSAv-AF555) that colocalized with a red signal (AF647-DNA-bt), *N*_*coloc* 22_, by the number of red signals (*N*_*total* 22_), and this value was corrected for the fraction of unoccupied DNA origami (Eq. 7).

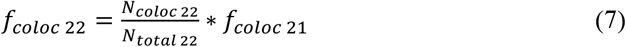

Finally, to determine the fraction of DNA origami functionalized with H57-scF_V_, DNA origami were pre-stained with YOYO and AS635P-conjugated H57-scF_*V*_ was used for two-color colocalization experiments. The number of red signals (H57-scF_*V*_) that colocalized with signals in the blue channel (YOYO), *N*_*coloc* 23_, divided by the number of blue signals (*N*_*total* 23_), yielded the fraction of DNA origami functionalized with H57-scF_V_ (Eq. 8).

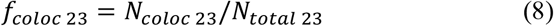

### Construct 3: H57-DNA

The second modification step (following incorporation of the handle) concerned the hybridization of a DNA-conjugated H57-scF_V_ to the handle. An AS635P-modified handle (DNA-AS635P) and AF555-labeled H57-DNA was applied to determine the fraction of DNA origami functionalized with DNA-conjugated H57-scF_V_ *(f*_*coloc 32*_), The number of green signals (H57-scF_*V*_) colocalizing with a red signal (DNA-AS635P), *N*_*coloc 32*_, was divided by the number of red signals (*N*_*total 32*_), and corrected for the fraction of DNA origami without handle, *f*_*coloc* 11_ (Eq. 9).

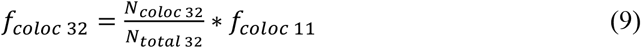

### Construct 4: H57-PNA

The fraction of DNA origami functionalized with PNA-conjugated H57-scF_V_ *(f*_*coloc* 42_) was determined in analogy to construct H57-DNA (Eq. 10).

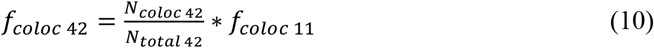

### Construct 5: H57-mSAv

The second modification step (following incorporation of the handle) was the hybridization of DNA-coupled mSAv to the handle. For determining the fraction of DNA origami functionalized with mSAv-DNA, *f*_*coloc* 52_, DNA origami were pre-stained with YOYO and AS635P-labeled mSAv-DNA was used. By determining the number of signals in the red color channel (mSAv-DNA-AS635P) that colocalized with a signal in the blue color channel (YOYO), *N*_*coloc* 52_, divided by the number of detected blue signals, *N*_*total* 52_, *f*_*coloc* 52_ could be derived via (Eq. 11).

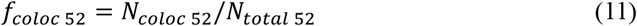

The fraction of DNA origami functionalized with H57-scF_V_, *f*_*coloc 53*_, was determined by using mSAv-DNA-AS635P and AF555-conjugated H57-scF_*V*_. Here, the number of green signals (H57-scF_V_) that colocalized with a red signal (mSAv-DNA-AS635P), *N*_*coloc 53*_, were divided by the number of red signals (*N*_*total 53*_) and corrected for the fraction of DNA origami without mSAv-DNA, yielding the fraction of DNA origami functionalized with H57-scF_V_, *f*_*coloc 53*_ (Eq. 12).

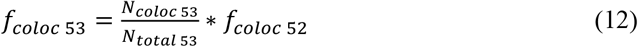

### Construct 6: H57-tSAv

The first two steps for generating this construct were analogous to those involved in the generation of construct 1: dSAv-H57.

The fraction of DNA origami functionalized with tSAv, *f*_*coloc 63*_, was determined by fluorescently labeling DNA origami with YOYO and using AS635P-labeled tSAv. The number of red signals (tSAv) that colocalized with a blue signal (YOYO), *N*_*coloc 63*_, was divided by the number of blue signals (*N*_*total 63*_). Thus, the fraction of DNA origami functionalized with tSAv is given by Eq. 13.

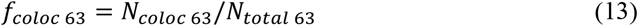

The fraction of DNA origami functionalized with H57-scF_V_, *f*_*coloc 64*_, was derived in analogy to Eq. 6 for H57-dSAv (Eq. 14).

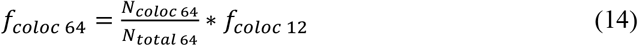

### Construct 7: H57-tSAv-NL

Construct H57-tSAv-NL was created in analogy to H57-dSAv-NL, yielding the fraction of biotin-bound tSAv via Eq. 15.

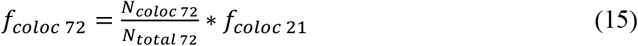

and the fraction of DNA origami functionalized with H57-scF_V_ via Eq. 16.

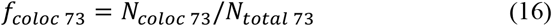

### Determination of functionalization stoichiometry via brightness analysis

Two-color colocalization analysis yielded the fraction of DNA origami carrying at least one H57-scF_V_. To determine the number of ligands on a functionalized DNA origami construct, we used single molecule brightness analysis based on a MATLAB (Mathworks)-based maximum-likelihood estimator to determine position, integrated brightness *B*, full width at half-maximum (FWHM) and local background of individual signals in the images as described previously ^36,37^. Briefly, functionalized DNA origami were anchored to SLBs, and the integrated brightness *B* was determined for all recorded positions. Images were taken at multiple different locations (*n* ≥ 10) yielding a minimum of ∼800 signals. The brightness values *B* of a monomer reference (a SLB-anchored single H57-scF_V_ molecule labeled with AF555) were used to calculate the probability density function (pdf) of monomers, *ρ*_1_(*B*). Because of the independent photon emission process, the corresponding pdfs of *N* colocalized emitters can be calculated by a series of convolution integrals:

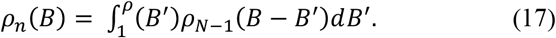

A weighted linear combination of these pdfs was used to calculate the brightness distribution of a mixed population of monomers and oligomers:

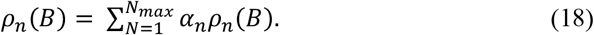

Brightness values for each DNA origami construct were pooled and used to calculate *ρ*(*B*). A least-square fit with Eq. 18 was employed to determine the weights of the individual pdfs, *α*_*N*_, with 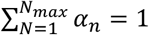. For fits of constructs 1 - 5, no higher contributions than monomers (*α*_1_) were observed.

For fits of construct 6 & 7, contributions up to trimers (*α*_*3*_) were observed.

### Mobility of DNA origami constructs on SLBs

For diffusion analysis of DNA origami constructs, at least 10 image sequences with 100 images each were recorded at different locations on the SLB at an illumination time of 3 ms and a time lag of 10 ms. Images were analyzed using in-house algorithms implemented in MATLAB^38^. Mean-square displacements (MSDs) were averaged over all trajectories, plotted as a function of time lag *t* and the diffusion coefficient *D* was determined by fitting the function *MSD* = 4*Dt* + 4*σ*_*xy*_, where *σ*_*xy*_ denotes the localization precision; diffusion coefficients were determined from the first two data points of the MSD-t-plot.

### Determination of H57-scF_*V*_ surface densities

Average surface densities of AF555-labeled H57-scF_*V*_ on SLBs were determined by dividing mean intensities per µm^2^ recorded at 8 different positions on the SLB by the brightness of a single AF555-H57-scF_V_ molecule.

### Determination of TCR surface densities

Average TCR surface densities were calculated from T-cells in contact with ICAM-1–functionalized SLBs and labeled to saturation with H57-scF_V_ variants (DNA-conjugated H57-scF_V_, PNA-conjugated H57-scF_V_, biotinylated H57-scF_V_, H57-scF_V_) fluorescently labeled with AF555 ^35^. T-cell brightness per µm^2^ was then divided by the brightness of a single AF555-H57-scF_V_molecule.

### Calcium imaging experiments and analysis

10^6^ T-cells were incubated in T-cell media supplemented with 5 µg ml^-1^ Fura-2 AM (11524766, ThermoFisher Scientific) for 20 min at 24°C. Excessive Fura-2 AM was removed by washing 3x with HBSS + 2% FBS. T cells were diluted with HBSS + 2% FBS to get a final concentration of 5*10^3^ cells µl^-1^. 10^5^ cells were transferred to the Lab-Tek™chamber and image acquisition was started immediately after T cells landed on the functionalized SLBs. Fura-2 AM was excited using a monochromatic light source (Polychrome V, TILL Photonics), coupled to a Zeiss Axiovert 200M equipped with a 10x objective (Olympus), 1.6x tube lens and an Andor iXon Ultra EMCCD camera. A longpass filter (T400lp, Chroma) and an emission filter were used (510/80ET, Chroma). Imaging was performed with excitation at 340 nm and 380 nm, with illumination times of 50 ms and 10 ms, respectively. The total recording time was 10 min at 1 Hz. Precise temperature control was enabled by an in-house-built incubator equipped with a heating unit. Calcium experiments were carried out at 37°C.

ImageJ was used to generate ratio and sum images of 340nm/380nm. T cells were segmented and tracked via the sum image of both channels using an in-house Matlab algorithm based on Gao et al. ^39^. Cellular positions and tracks were stored and used for intensity extraction based on the ratio image. Intensity traces were normalized to the starting value at time point zero. Traces were categorized in “activating” and “non-activating” based on an activation threshold ratio of 0.4. The activation threshold was chosen based on comparison of individual traces of a positive control (ICAM-1 100 µm^-2^, B7-1 100 µm^-2^, pMHC 150 µm^-2^) and a negative control (ICAM-1 100 µm^-2^, B7-1 100 µm^-2^) (*n >* 40). For generating dose-response curves, at least 15 calcium measurements (of typically ∼100 cells in a region of interest) were conducted, with each measurement at a specific ligand density. The percentage of activated cells was evaluated for each measurement and normalized to the positive control. Data were plotted as % activated cells *A* as a function of ligand surface densities *L* to generate dose-response curves and fitted with Eq. 19 to extract the activation threshold *T*_*A*_, the maximum response *A*_*max*_ and the Hill coefficient *n*.

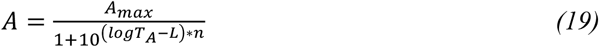

All fit parameters and their 95% confidence intervals (CI) are summarized in the **SI Table S3**. Statistical significance between the values *T*_*A*,1_ and *T*_*A*,2_for two different data sets was determined via a bootstrap ratio test ^40,41^ as follows: A bootstrap sample was obtained by drawing *n* data points (sampling with replacement) from a dose-response curve, where *n* equals the size of the data set. From each data set, 1000 bootstrap samples were drawn and fitted via Eq. 19. This yielded threshold values 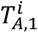 and 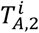 *(i* = 1, …, 1000) for each of the bootstrap samples from the two different data sets. The ratio 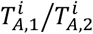 (*i* = 1, …, 1000) was calculated for each pair of bootstrap samples. If the 100 * (1 *- α)*% CI of 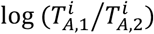 did not contain 0, the null hypothesis of equal *T*_*A*_ was rejected at a significance level of *α*.

### Protein expression, purification and conjugation

The TCRβ-reactive H57 single chain antibody fragment (H57-scF_V_) featuring an unpaired cysteine at the C-terminus (S248C) and the H57-scF_*V*_ equipped with a C-terminal BirA ligase biotinylation site were prepared as described ^42,6^. Both H57-scF_V_ versions were cloned into a pET21a(+) expression vector for expression in *Escherichia coli* (BL-21). Insoluble inclusion bodies were extracted via sonication and carefully washed with 1% Triton (Merck) and 1 % deoxycholic acid (Merck) in 50 mM Tris pH 8.0, 0.2 M NaCl and 2mM EDTA (all Merck) before dissolving them finally in 6 M guanidine hydrochloride (Merck). H57-scF_V_ were refolded from inclusion bodies by a stepwise reduction of the guanidine hydrochloride concentration within the refolding buffer (50 mM Tris pH 8.0, 0.2 M NaCl, 1 mM EDTA) and shifting the redox system from reducing to oxidizing conditions ^43^. After a final dialyzing step against 1x PBS, refolded H57-scF_V_ were concentrated using Amicon®Ultra-15 centrifugal filters with a 10 kDa cutoff (Merck) and purified via gel filtration using Superdex 200 (10/300, Cytiva) on an Äkta pure chromatography system (Cytiva). H57 scF_V_ containing an unpaired cysteine were concentrated in the presence of 50 μM Tris-(2-carboxyethyl) phosphine hydrochloride (TCEP, Pierce).

Monomeric H57-scF_V_ featuring a BirA recognition site was site-specifically biotinylated using a biotin ligase (Avidity) at 30 °C, followed by a buffer exchange to 1x PBS via gel filtration (Superdex-75, 30/300 Cytiva). Biotinylated H57-scF_V_ were randomly conjugated on surface-exposed lysines with Alexa Fluor 555 (AF555) carboxylic acid, succinimidyl ester (ThermoFisher Scientific) or Abberior Star 635P (AS635P) carboxylic acid, succinimidyl ester (Abberior) according to the manufacturer’s instructions. To remove excess dye, the AF555- or AS635P-conjugated and biotinylated H57-scF_*V*_ were purified via gel filtration using Superdex 75 (10/300 GL, Cytiva). Fractions containing monomeric, fluorescently-labeled and biotinylated H57-scF_*V*_ were concentrated to 0.2 - 1 mg/mL with 10 kDa Amicon®Ultra-4 centrifugal filters (Merck) and stored in 1x PBS supplemented with 50 % glycerol at -20°C. The protein to dye ratio ranged between 0.93 and 1.1 for the AF555-labeled H57-scF_*V*_ and was 2.0 for the AS635P-labeled H57-scF_*V*_ as determined by spectrophotometry (280 to 555 nm or 280 to 638 nm ratio).

H57-scF_V_ featuring a free cysteine at the C-terminus were conjugated to dibenzylcyclooctyne-maleimide (DBCO-maleimide, Jena Bioscience) in the presence of 50 µM TCEP for 2 h at room temperature followed by a gel filtration step (Superdex 75, 30/300 Cytiva) to remove unreacted DBCO-maleimide. Directly thereafter, monomeric H57-scF_V_-DBCO was labeled with AF555 carboxylic acid, succinimidyl ester (ThermoFisher Scientific) and purified via gel filtration to remove excess unconjugated dye. In the last step, AF555-conjugated H57-scF_V_-DBCO was coupled to Azido-PEG4-DNA (TTTTACATGACACTACTCCAC) or Azido-PNA (see below) for 2.5 h at room temperature, purified via gel filtration (Superdex 75, 30/300 Cytiva) to remove unreacted Azido-PEG4-DNA or Azido-PNA, concentrated with 10 kDa Amicon®Ultra-4 centrifugal filters (Merck) and stored in 1x PBS supplemented with 50% glycerol at -20°C. The protein to AF555-dye ratio was 1.1 for the H57-DNA, and 1.15 for the H57-PNA as determined by spectrophotometry (280 to 555 nm ratio) before conjugation to Azido-PEG4-DNA or Azido-PNA. To arrive at Azido-PNA, we functionalized PNA-cysteine (O - TTACATGACACTACTCCAC) with Azido-PEG3-maleimide (Jena Bioscience) according to the manufacturer’s instructions (Azido-PEG3-Maleimide Preparation Kit). The product was purified via reversed phase chromatography (1260 Infinity II, Agilent Technologies) on a C18 column (Pursuit XRs 5 C18 250 × 21.2 mm) to separate PNA-cysteine from Azido-PNA. Positive fractions containing only Azido-PNA were verified by MALDI-TOF mass spectrometry (Bruker).

Monovalent streptavidin (mSAv) featuring an unpaired cysteine (A106C) in the biotin-binding subunit was produced as described^44^. After refolding from inclusion bodies and purification via anion exchange chromatography (Mono Q, 5/50, Cytiva) and gel filtration (Superdex 200, 30/300, Cytiva), mSAv was labeled randomly on lysine residues with AS635P carboxylic acid, succinimidyl ester (Abberior) according to the manufacturer’s instructions and purified via gel filtration (Superdex 200, 30/300 Cytiva). Fractions containing STAR635P-conjugated mSAV were concentrated with 10 kDa Amicon®Ultra-4 centrifugal filters (Merck) and conjugated to trans-Cyclooctene-PEG3-Maleimide (TCO-PEG3-Maleimide, Jena Bioscience) and again purified via gel filtration (Superdex 200, 30/300, Cytiva) to remove unreacted TCO-PEG3-Maleimide. Finally, AS635P-labeled mSAv-TCO was conjugated to tetrazine-PEG5-oligo (TTTTACATGACACTACTCCAC) for 2h at room temperature and purified via gel filtration (Superdex 75, 30/300, Cytiva). Monomeric STAR635P-labeled mSAv-DNA was concentrated with 10 kDa Amicon®Ultra-4 centrifugal filters (Merck) and stored in 1x PBS supplemented with 50 % glycerol at -20 °C. The protein to dye ratio for the AS635P-labeled mSAv-DNA was 1.0 as determined by spectrophotometry (280 to 638 nm ratio) before conjugation with TCO-PEG3-Maleimide.

Trans-divalent streptavidin (dSAv) and tetravalent streptavidin (tSAv) were prepared based on a protocol by Fairhead *et al*. ^24^ and as described in ^6^.

Tetravalent streptavidin was refolded from inclusion bodies containing only biotin-binding streptavidin subunits and purified via gel filtration (Superdex 200, 30/300, Cytiva). Monomeric fractions were labeled with Abberior Star 635P (AS635P) carboxylic acid, succinimidyl ester (Abberior) according to the manufacturer’s instructions and purified via gel filtration (Superdex 200, 30/300 Cytiva). Fractions containing STAR635P-conjugated mSAv were concentrated with 10 kDa Amicon®Ultra-4 centrifugal filters (Merck) and stored in 1x PBS supplemented with 50 % glycerol at -20 °C. The protein to dye ratio for the AS635P-labeled tSAv was 1.2 as determined by spectrophotometry (280 to 638 nm ratio). 2xHis_6_-tag pMHC-AF555 was produced as described in ^35^.

### Tissue culture

Primary T-cells isolated from lymph nodes or spleen of 5c.c7 *αβ* TCR transgenic mice were pulsed with 0.5 µM moth cytochrome c peptide (MCC) 88-103 peptide (C18-reverse phase HPLC-purified; sequence: ANERADL<u>IAYLKQATK</u>, T-cell epitope underlined, Elim Biopharmaceuticals Inc, USA) and 50 U ml^-1^ IL-2 (eBioscience) for 7 days to expand CD4+ T-cells and arrive at an antigen-experienced T-cell culture ^45^. T cells were maintained at 37°C and 5% CO_2_ in RPMI 1640 media (Life technologies) supplemented with 10% FBS (MERCK), 100 µg ml^-1^ penicillin (Life technologies), 100 µg ml^-1^ streptomycin (Life technologies), 2 mM L-glutamine (Life technologies), 0.1 mM non-essential amino acids (Lonza), 1 mM sodium pyruvate (Life technologies) and 50 µM *β*-mercaptoethanol (Life technologies). Dead cells were removed six days after T-cell isolation with a density-dependent gradient centrifugation step (Histopaque 1119, Sigma). Antigen-experienced T-cells were used for experiments on day 7 – 9.

### Animal model and ethical compliance statement

5c.c7 αβ TCR-transgenic mice bred onto the B10.A background were a kind gift from Michael Dustin (University of Oxford,UK). Both male and female mice at 8-12 weeks old were randomly selected and sacrificed for isolation of T-cells from lymph nodes and spleen, which was evaluated by the ethics committees of the Medical University of Vienna and approved by the Federal Ministry of Science, Research and Economy, BMWFW (BMWFW-66.009/0378-WF/V/3b/2016). Animal husbandry, breeding and sacrifice of mice was performed in accordance to Austrian law (Federal Ministry for Science and Research, Vienna, Austria), the guidelines of the ethics committees of the Medical University of Vienna and the guidelines of the Federation of Laboratory Animal Science Associations (FELASA), which match those of Animal Research: Reporting in vivo Experiments (ARRIVE). Further, animal husbandry, breeding and sacrifice for T-cell isolation was conducted under Project License (I4BD9B9A8L) which was evaluated by the Animal Welfare and Ethical Review Body of the University of Oxford and approved by the Secretary of State of the UK Home Department. They were performed in accordance to Animals (Scientific Procedures) Act 1986, the guidelines of the ethics committees of the Medical Science of University of Oxford and the guidelines of the Federation of Laboratory Animal Science Associations (FELASA), which match those of Animal Research: Reporting in vivo Experiments (ARRIVE).

## Acknowledgments

This work was supported by the Austrian Science Fund (FWF projects V538-B26 (ES), T134040-2010 (JH) and F6809-N36 (GS, MS); the PhD program Cell Communication in Health and Disease W1205, RP, JBH and HS), the TU Wien doctoral college BioInterface (JH), the Vienna Science and Technology Fund (WWTF, LS13-030, GJS and JBH), the Boehringer Ingelheim Fonds (RP), the Wellcome Trust (Principal Research Fellowship 100262 Z/12/Z, EK) and the Kennedy Trust for Rheumatology Research (EK).

