## Supplementary Information for "Strategies for the site-specific decoration of DNA origami nanostructures with functionally intact proteins"

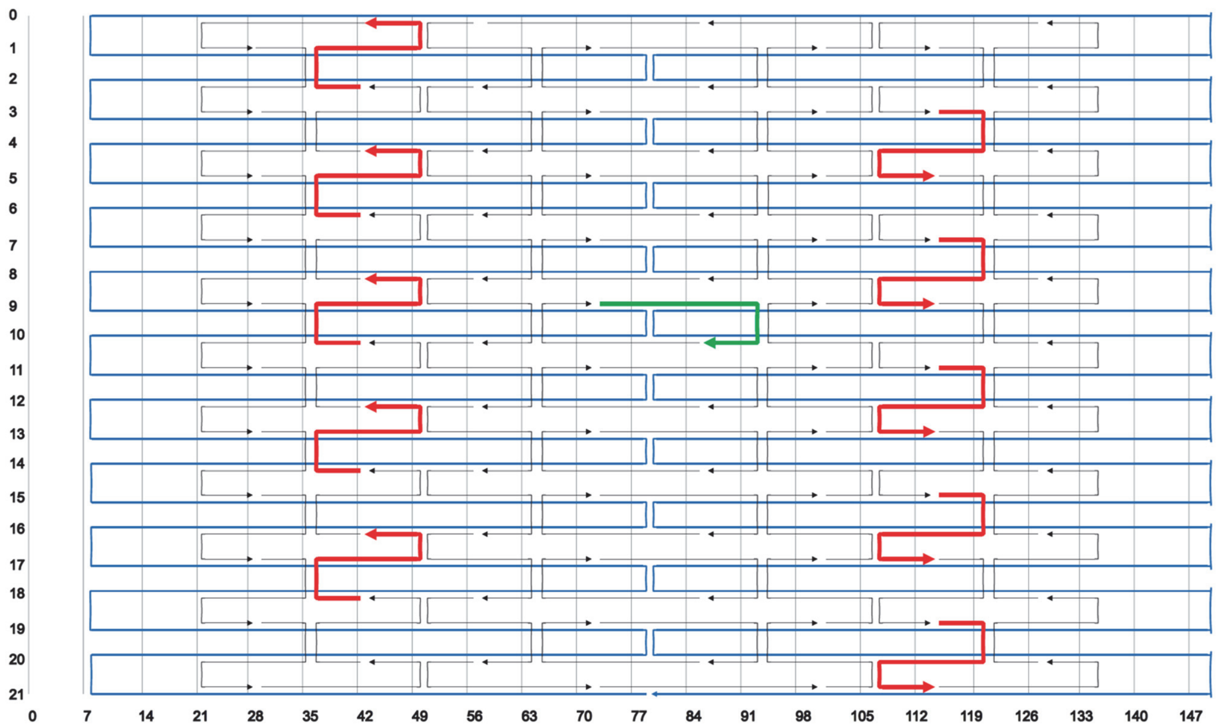


**Fig. S1. DNA origami scaffold routing.** The DNA origami rectangular tile (65x54 nm) was designed based on the M13mp18 scaffold[1] using caDNAno[2]. At the site chosen for ligand attachment, the staple strand was elongated at its 3'-end with 21 bases (green arrow). For attachment to the SLB via complementary cholesterol-oligonucleotides, staple strands were elongated at predefined positions at their 5'-end with 25 bases as indicated by the red arrows.

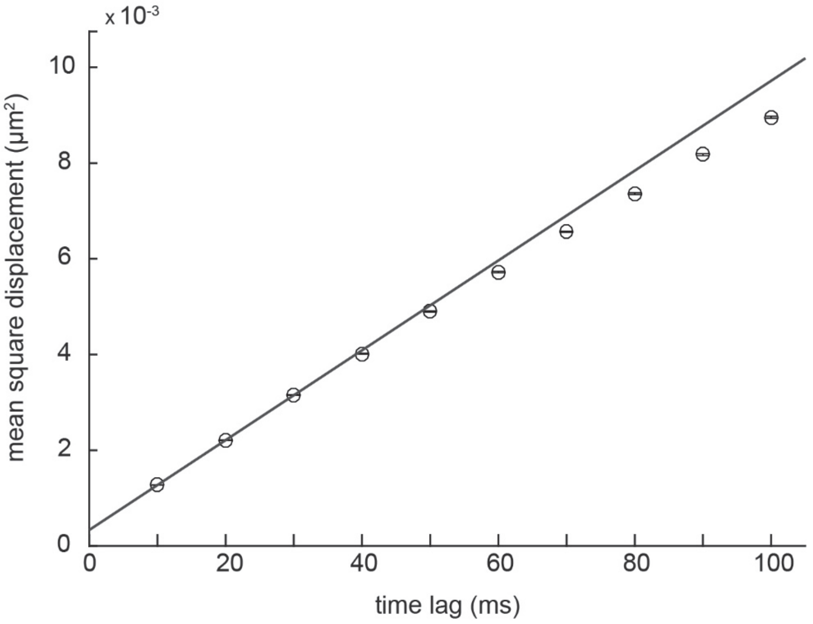


**Fig. S2. Mobility of functionalized DNA origami structures on fluid-phase SLBs.** Individual DNA origami structures were tracked on POPC SLBs at a frame rate of 100 Hz. Mean square displacements were determined and plotted as a function of time lags. Assuming free Brownian motion, the diffusion coefficient D was derived by fitting the first two data points with a linear fit. An exemplary plot for H57-dSAv is shown.

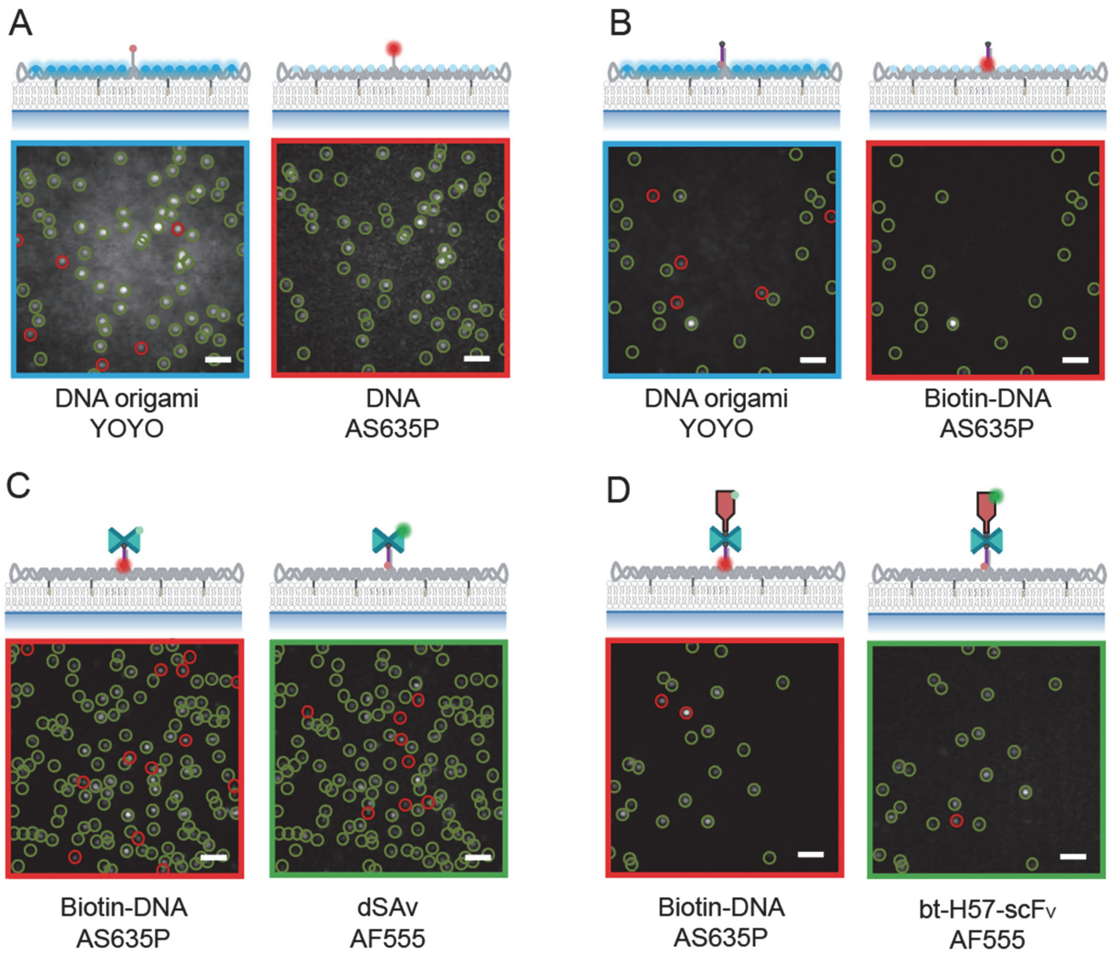


**Fig. S3. Determining origami conjugation efficiencies.** Single molecule two-color co-localization imaging was applied to determine the efficiency of each modification step. *(A)* First, we evaluated the availability of elongated staple strands (handles) for site-specific hybridization. For this purpose, we used fluorophore-conjugated handles (DNA-AS635P) and labeled the DNA origami structure with YOYO, a DNA-intercalating fluorophore. The percentage of co-localized signals in the blue (YOYO) and the red (DNA-AS635P) color channel yielded the handle incorporation efficiency. Representative TIRF images of DNA origami on an SLB are shown. Green open circles indicate signals detected in both color channels; red open circles indicate signals detected only in one channel. *(B)* Two-color colocalization of fluorescently labeled biotinylated oligo nucleotides (bt-DNA-AS635P) and YOYO yielded the efficiency of hybridization to the handle. *(C)* Binding of dSAv to the biotinylated handle was then determined via colocalization of fluorescently labeled biotinylated oligos (bt-DNA-AS635P) and dSAv (dSAv-AF555). *(D)* Finally, two-color colocalization of hybridized bt-DNA-AS635P and site-specifically biotinylated AF555-labeled H57-scFvs yielded the overall functionalization efficiency of DNA origami with the TCR-ligand. The functionalization efficiency was then assessed as described in the methods section. Scale bar, 2 µm.

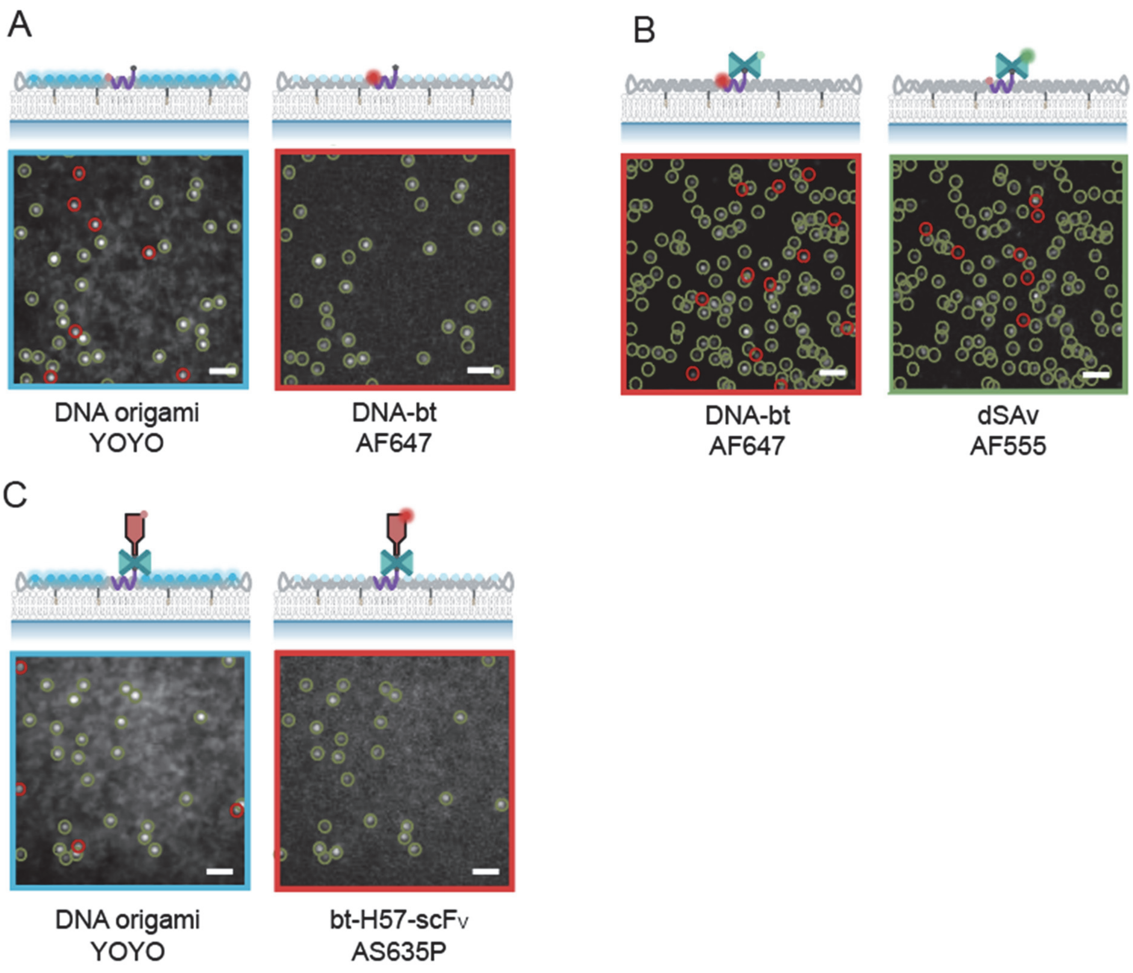


**Fig. S4. Functionalization efficiencies determined for the construct H57-dSAv-NL.** *(A)* To determine the incorporation efficiency of the biotinylated handle, DNA origami structures featuring a fluorescently labeled, biotinylated handle (AF647-DNA-bt) were produced and unspecifically stained with the DNA-intercalating dye YOYO. Representative TIRF images of DNA origami on an SLB are shown. Green open circles indicate signals detected in both color channels; red open circles designate signals detected only in one channel. The percentage of co-localized signals in the blue (YOYO) and red (AF647-DNA-bt) color channel yielded the incorporation efficiency. *(B)* The binding efficiency of dSAv to the biotinylated handle was determined via two-color colocalization of fluorescently labeled dSAv (dSAv-AF555) and AF647-DNA-bt. *(C)* Two-color colocalization of AS635P-labeled H57-scFvs and YOYO-stained DNA origami structures yielded the overall functionalization efficiency of DNA origami with the TCR-ligand. Scale bar, 2 µm.

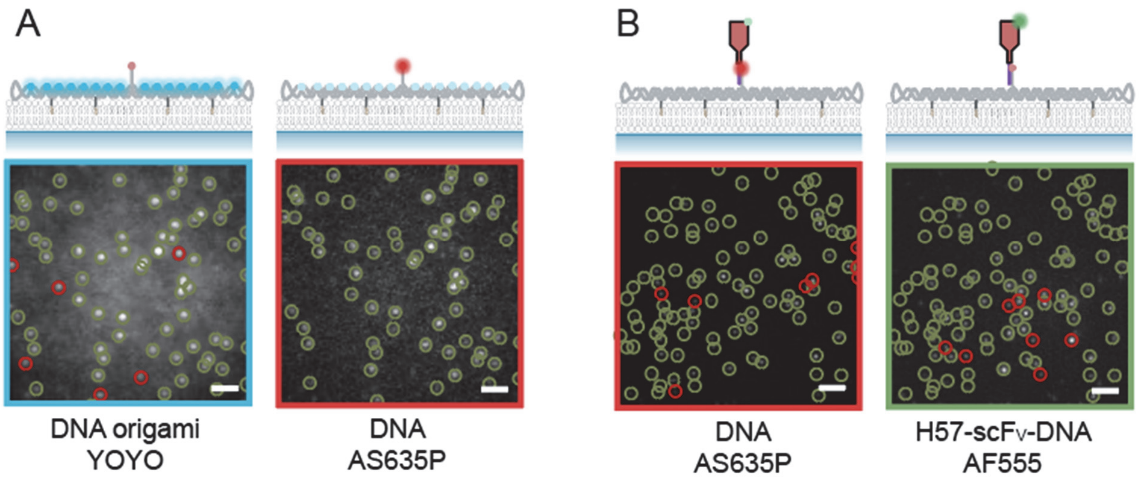


**Fig. S5. Determining functionalization efficiencies for construct H57-DNA.** *(A)* Handle availability was determined via two-color co-localization of DNA-AS635P and YOYO. Representative TIRF images of DNA origami on a SLB are shown. Green open circles indicate signals detected in both color channels; red open circles designate signals detected only in one channel. *(B)* Colocalization of AF555-labeled DNA-conjugated H57-scFvs (H57-DNA) and DNA-AS635P yielded the overall functionalization efficiency of DNA origami with ligand. Scale bar, 2 μm.

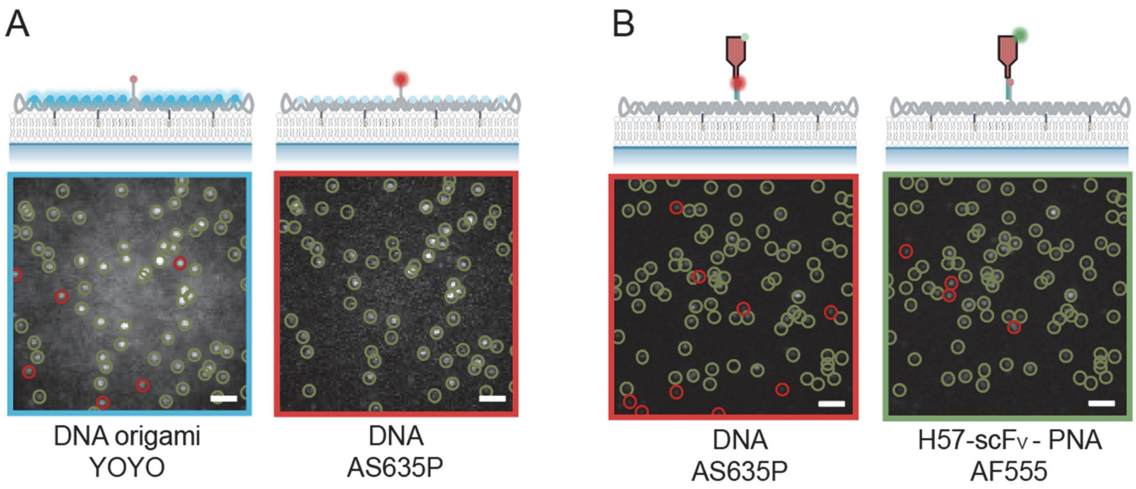


**Fig. S6. Determining functionalization efficiencies for the construct H57-PNA.** *(A)* Handle availability was determined via two-color co-localization of DNA-AS635P and YOYO. Exemplary TIRF images of DNA origami on a SLB are shown. Green open circles indicate signals detected in both color channels; red open circles designate signals detected only in one channel. *(B)* Colocalization of AF555-labeled PNA-conjugated H57-scFVs (H57-PNA) and DNA-AS635P yielded the overall functionalization efficiency of DNA origami with ligand. Scale bar, 2 µm.

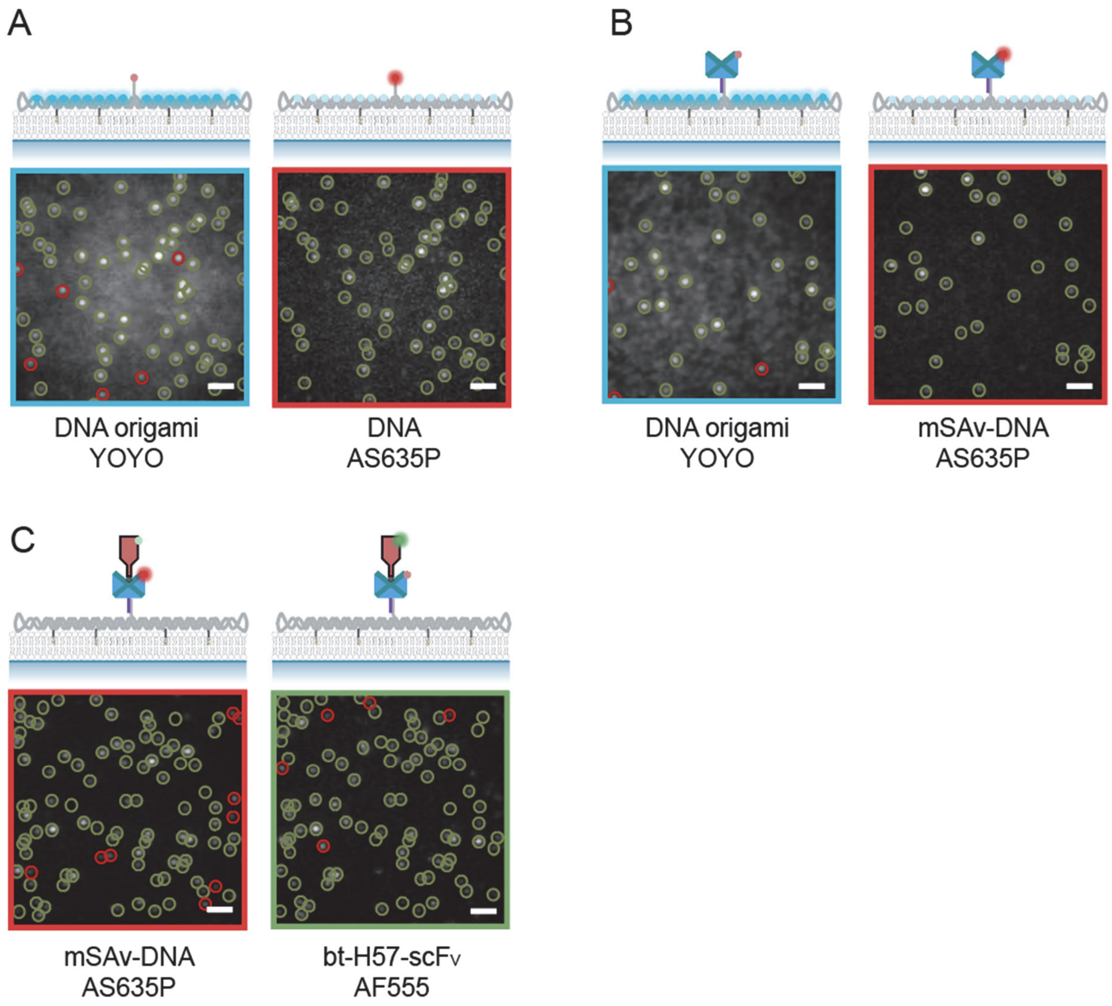


**Fig. S7. Determining functionalization efficiencies for the construct H57-mSAv.** *(A)* Handle availability was determined by two-color co-localization of DNA-AS635P and YOYO. Exemplary TIRF images of DNA origami on a SLB are shown. Green open circles indicate signals detected in both color channels; red open circles indicate signals detected only in one channel. *(B)* Colocalization of AS635P-labeled DNA-coupled mSAv (mSAv-DNA-AS635P) and YOYO yielded functionalization efficiency with mSAv. *(C)* Finally, two-color colocalization of mSAv-DNA-AS635P and AF555-labeled biotinylated H57-scFvs yielded the overall functionalization efficiency of DNA origami with ligand. Scale bar, 2 µm.

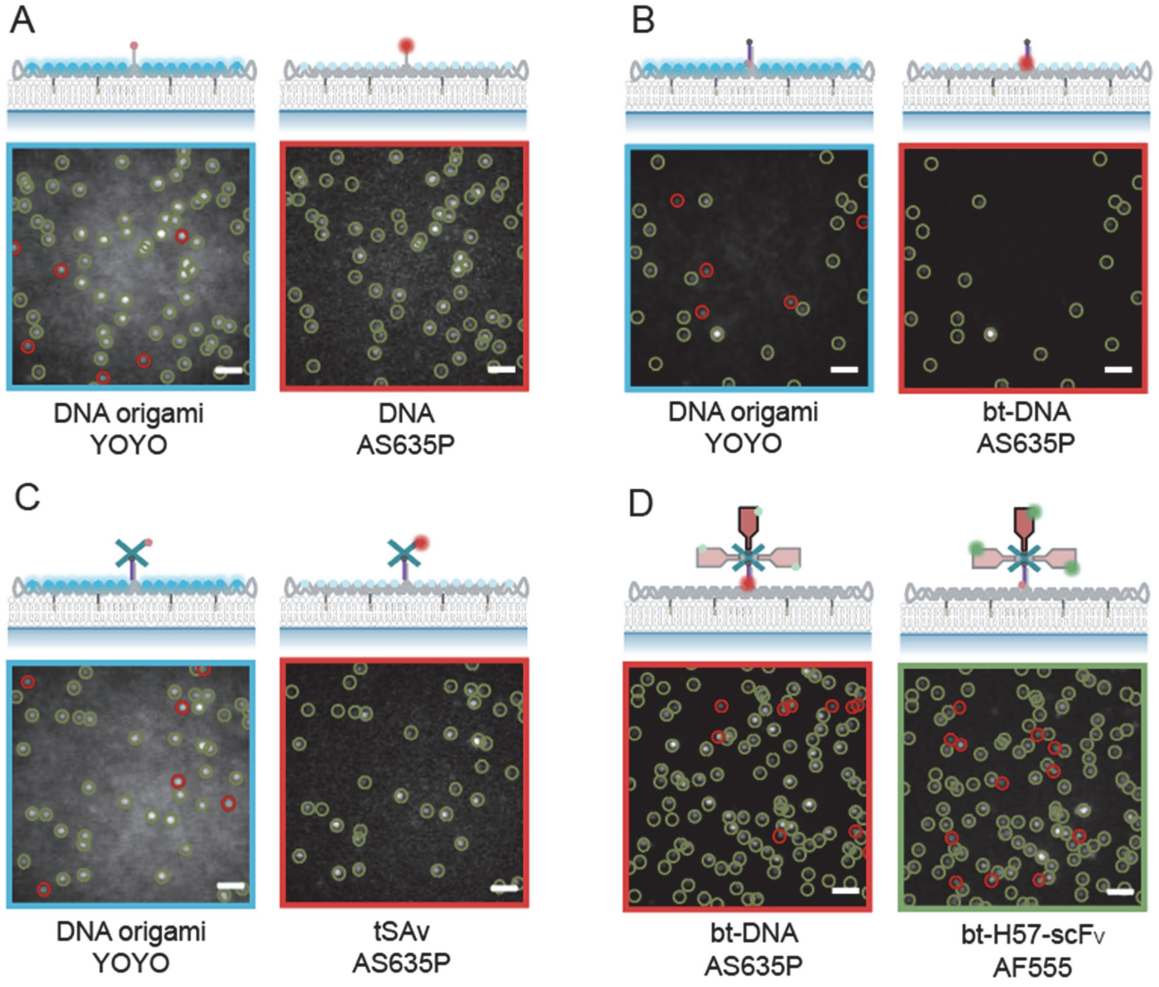


**Fig. S8. Determining functionalization efficiencies for construct H57-tSAv.** *(A)* Handle availability was determined by two-color co-localization of DNA-AS635P and YOYO. Exemplary TIRF images of DNA origami on a SLB are shown. Green open circles indicate signals detected in both color channels; red open circles indicate signals detected only in one channel. *(B)* Next, two-color colocalization of fluorescently labeled biotinylated oligos (bt-DNA-AS635P) with YOYO yielded the efficiency of hybridization of biotinylated oligo to the handle. *(C)* Colocalization of fluorescently labeled tSAv (tSAv-AS635P) and YOYO yielded functionalization efficiency with tSAv. *(D)* Finally, colocalization of hybridized biotin-DNA-AS635P and AF555-labeled H57-scFvs yielded the overall functionalization efficiency of DNA origami with ligand. Scale bar, 2 µm.

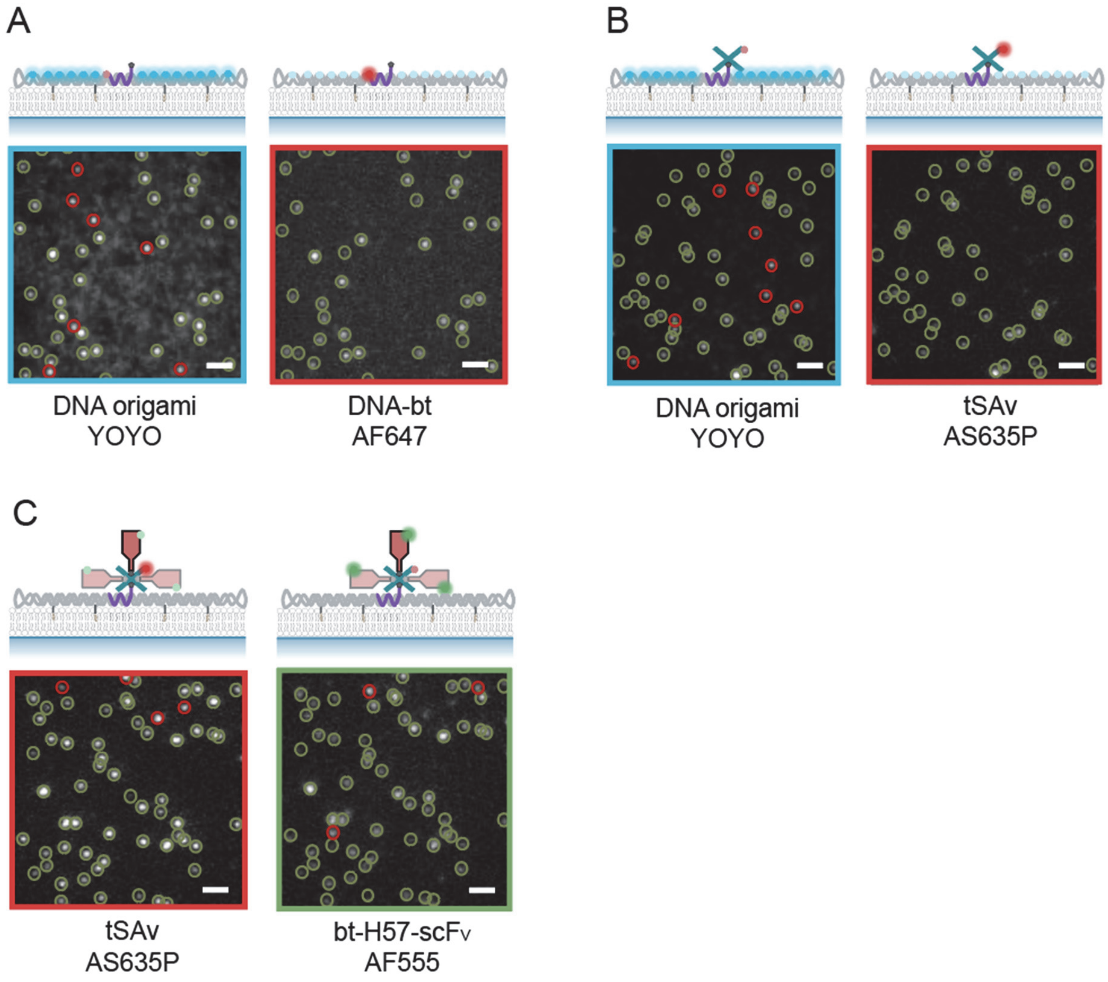


**Fig. S9. Determining functionalization efficiencies for construct H57-tSAv-NL.** *(A)* Incorporation of the biotinylated handle on DNA origami for functionalization was assessed via colocalization of a fluorescently labeled biotinylated oligo (AF647-DNA-bt) and YOYO. Exemplary TIRF images of DNA origami on a SLB are shown. Green open circles indicate signals detected in both color channels; red open circles indicate signals detected only in one channel. *(B)* The binding efficiency of tSAv to the biotinylated handle was determined via two-color colocalization of fluorescently labeled tSAv (tSAv-AS635P) and YOYO. *(C)* Last, two-color colocalization of tSAv-AS635P and AF555-labeled H57-scFvs yielded the overall functionalization efficiency of DNA origami with ligand. Scale bar, 2 μm.

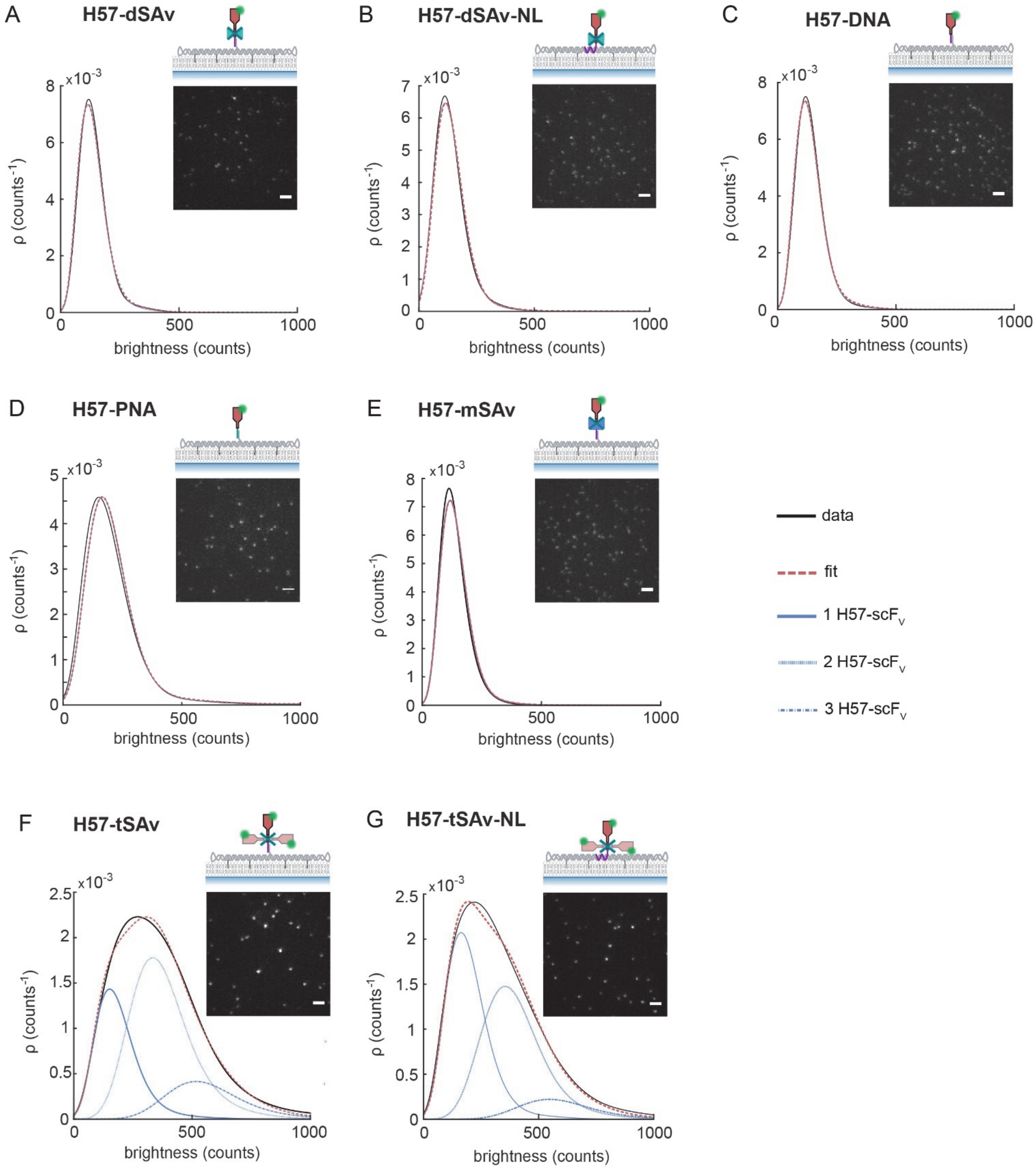


**Fig. S10. Determining functionalization stoichiometries for H57-scFV-functionalized DNA origami constructs.** Exemplary TIRF images and corresponding brightness distributions ρ of DNA origami constructs functionalized with AF555-labeled H57-scFV on SLBs. The detected signals were fitted and deconvolved into monomer and multimer contributions[3] (see methods section). Scale bar, 2 μm.

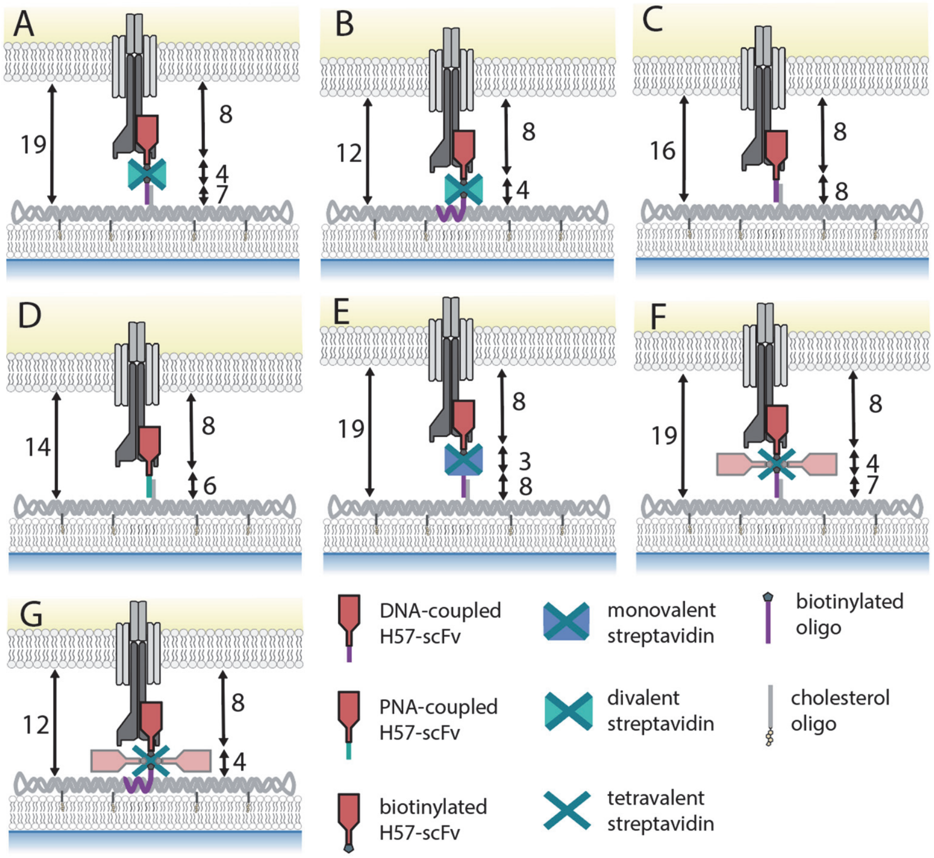


**Fig. S11. Axial dimensions at the T-cell – SLB interface.** Schematic sketches of TCR engagement for the different ligand-functionalized DNA origami structures: H57-dSAv *(A)*, H57-dSAv-NL *(B)*, H57-DNA *(C)*, H57-PNA *(D)*, H57-mSAv *(E)*, H57-tSAv *(F)*, H57-tSAv-NL *(G)*. Distances were estimated from the protein crystal structures[5–8] and are given in nm.

**Note on length estimates**

The two *trans* biotin binding sites in SAv are separated by ~3.5 nm[8]; hence the contribution to the total length of the construct was assumed with 4 nm *(A, F)*. For mSAv *(E)*, this contribution was estimated with 3 nm from the crystal structure[7].

The DNA linker consists of 17 paired bases in constructs A, C, E, F; with the following variations:

**A, F:** (constructs H57-dSAv and H57-tSAv): 4 unpaired Ts on the handle, 17 paired bases, TEG linker and biotin. (~ 7 nm)

**C, E:** (constructs H57-DNA, H57-mSAv)): 4 unpaired Ts on the handle, 17 paired bases and 4 unpaired Ts on the hybridized DNA oligo, TEG linker (~ 8 nm)

**D:** The DNA/PNA linker consists of 4 unpaired Ts on the handle, 17 paired bases, 2 unpaired Ts and an O-linker (~ 6 nm).

**B, G:** (constructs H57-dSAv-NL and H57-tSAv-NL): The biotin is attached to the handle via 2 unpaired Ts and a TEG linker.

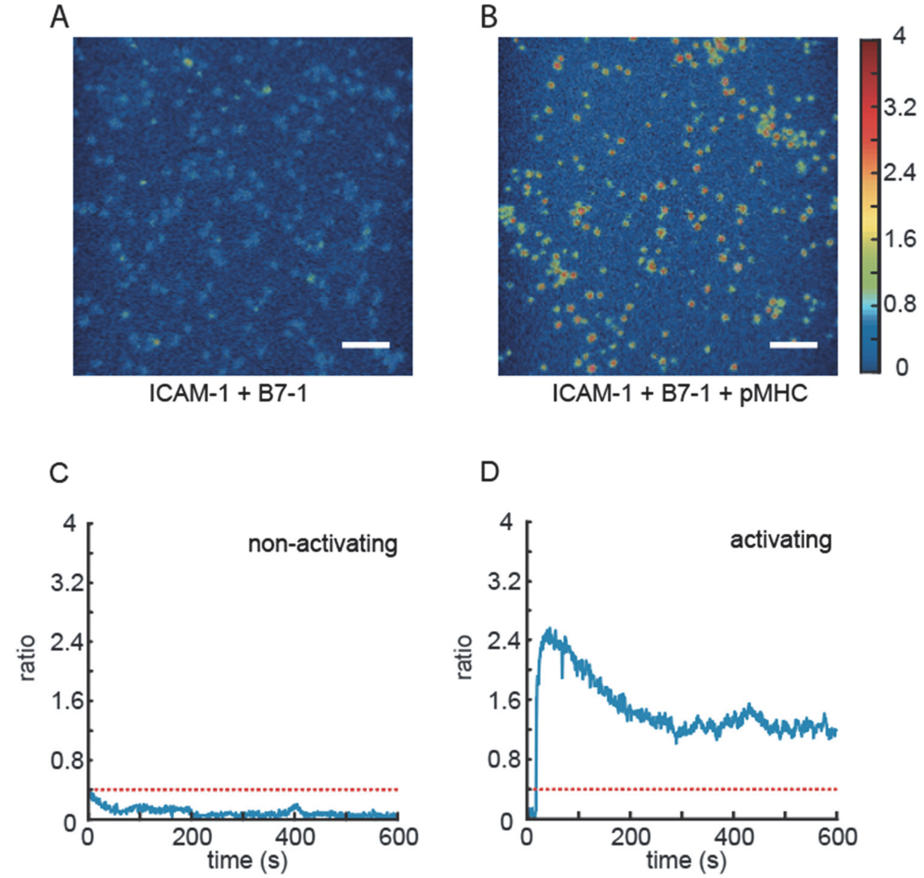


**Fig. S12. Calcium imaging experiments to assess the T-cell activation state.** T-cells were loaded with the ratiometric $Ca^{2+}$-sensitive dye Fura-2 AM, seeded onto SLBs and fluorescence emission was recorded at excitation wavelengths 340 nm and 380 nm over 10 min. Activation was tracked via a change of the intensity ratio (340/380nm). Exemplary ratio images recorded at activating (ICAM-1 100 $\mu m^{-2}$, B7-1 100 $\mu m^{-2}$, pMHC 150 $\mu m^{-2}$, *(A))* and non-activating (ICAM-1 100 $\mu m^{-2}$, B7-1 100 $\mu m^{-2}$, *(B))* conditions at 37°C are shown 5 min after cell seeding. Scale bar, 4 µm. *(C,D)* For each cell, the intensity ratio 340/380nm was determined and plotted over time. Exemplary calcium traces for a T-cell under non-activating *(C)* and activating *(D)* conditions are shown. The threshold ratio for counting a cell as "activated" was set to 0.4 for all experiments, indicated by a red dashed line.

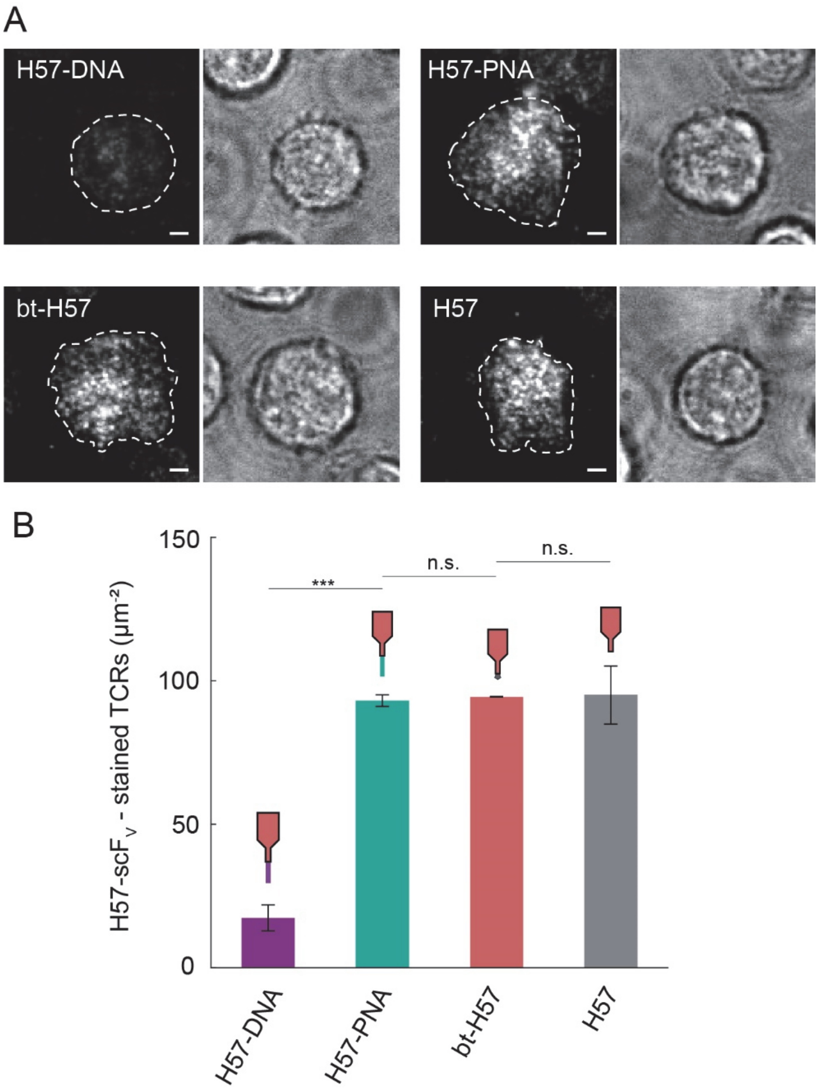


**Fig. S13. Determination of the TCR labeling efficiency of the different H57-scFv variants.** *(A)* TCRs on T-cells were stained with different AF555-conjugated H57-scFv variants (H57-DNA, H57-PNA, bt-H57, H57) and allowed to adhere to SLBs presenting 100 molecules per µm$^2$ ICAM-1 for imaging in TIRF microscopy mode. T-cells were labeled under saturating conditions for AF555-conjugated H57-scFv ( 20 µg ml$^{-1}$) [4]. The cell outline is indicated by a dashed white contour line. Images were recorded 5 -10 min after cell seeding. Scale bar, 2 µm. (B) Surface densities of labeled TCRs were quantified ($n \geq 27$ cells). Data are the mean of two independent experiments and two different mice (± s.e.m.).

**Table S1. Diffusion coefficients of constructs on SLBs.** Single-molecule trajectories of DNA origami structures functionalized with H57-scF$_V$ were recorded on SLBs, pooled and diffusion coefficients were determined by mean square displacement analysis. Diffusion coefficients are given as mean ± s.e.m. Data are from at least two independent experiments.

| Construct | D ($\mu m^2/s$) | trajectories (n) |
|---|---|---|
| **H57-dSAv** | 0.380 ± 0.013 | 8,203 |
| **H57-dSAv-NL** | 0.381 ± 0.015 | 9,857 |
| **H57-DNA** | 0.383 ± 0.012 | 7,601 |
| **H57-PNA** | 0.421 ± 0.003 | 3,797 |
| **H57-mSAv** | 0.385 ± 0.012 | 8,669 |
| **H57-tSAv** | 0.355 ± 0.015 | 11,108 |
| **H57-tSAv-NL** | 0.399 ± 0.001 | 5,699 |

**Table S2. Fluorophore pairs used for two-color colocalization microscopy.**

| Construct | Fluorophore ID | Handle incorporation | bt-DNA hybridization | SAv attachment | POI attachment |
|---|---|---|---|---|---|
| **1: H57-dSAv** | DYE 1 | DNA origami (YOYO) | DNA origami (YOYO) | bt-DNA (AS635P) | bt-DNA (AS635P) |
| | DYE 2 | DNA (AS635P) | bt-DNA (AS635P) | dSAv (AF555) | bt-H57 (AF555) |
| **2: H57-dSAv-NL** | DYE 1 | DNA origami (YOYO) | --- | DNA (AS635P) | DNA origami (YOYO) |
| | DYE 2 | DNA-bt (AF647) | --- | dSAv (AF555) | bt-H57 (AS635P) |
| **3: H57-DNA** | DYE 1 | DNA origami (YOYO) | --- | --- | DNA (AS635P) |
| | DYE 2 | DNA (AS635P) | --- | --- | H57-DNA (AF555) |
| **4: H57-PNA** | DYE 1 | DNA origami (YOYO) | --- | --- | DNA (AS635P) |
| | DYE 2 | DNA (AS635P) | --- | --- | H57-PNA (AF555) |
| **5: H57-mSAv** | DYE 1 | DNA origami (YOYO) | --- | DNA origami (YOYO) | mSAv-DNA (AS635P) |
| | DYE 2 | DNA (AS635P) | --- | mSAv-DNA (AS635P) | bt-H57 (AF555) |
| **6: H57-tSAv** | DYE 1 | DNA origami (YOYO) | DNA origami (YOYO) | DNA origami (YOYO) | bt-DNA (AS635P) |
| | DYE 2 | DNA (AS635P) | bt-DNA (AS635P) | tSAv (AS635P) | bt-H57 (AF555) |
| **7: H57-tSAv-NL** | DYE 1 | DNA origami (YOYO) | --- | DNA origami (YOYO) | tSAv (AS635P) |
| | DYE 2 | DNA-bt (AF647) | --- | tSAv (AS635P) | bt-H57 (AF555) |

**Table S3. Optimization of functionalization conditions for construct H57-dSAv.** For each step, the yield of functionalized DNA origami construct was determined for different conditions (incubation times and molar ratios) via two-color colocalization microscopy as sketched in SI Fig. S3. Optimal conditions for each step are marked in red; these were then used as the basis for subsequent steps. Cumulative yields are shown. Data are the mean (± s.e.m.) of at least two independent experiments.

| **Handle incorporation** | | | | | | |
|---|---|---|---|---|---|---|
| **Molar ratio DNA origami : handle oligo** | | | | | | |
| **1 : 1** | **1 : 2** | **1 : 5** | **1 : 10** | **1 : 30** | **1 : 50** | **1 : 100** |
| **19.1 ± 0.9** | **32.8 ± 0.8** | **46.6 ± 0.6** | **84.4 ± 2.1** | **83.4 ± 2.7** | **86.4 ± 0.3** | **86.4 ± 0.0** |

| **bt-DNA hybridization** | | | | | |
|---|---|---|---|---|---|
| **Molar ratio DNA origami : bt-DNA** | | | | | |
| **1 : 10** | **1 : 20** | **1 : 50** | **1 : 100** | **1 : 300** | **1 : 1000** |
| **26.9 ± 0.3** | **39.1 ± 2.0** | **86.4 ± 0.5** | **85.4 ± 0.3** | **85.4 ± 0.2** | **85.9 ± 0.0** |

| **SAv attachment** | | | | |
|---|---|---|---|---|
| **Molar ratio DNA origami : dSAv** | | | | |
| **1 : 1** | **1 : 5** | **1 : 10** | **1 : 100** | **Incubation time [min]** |
| **33.1 ± 2.4** | **34.1 ± 0.6** | **32.6 ± 1.5** | **67.1 ± 0.6** | **10** |
| **34.3 ± 1.7** | **61.0 ± 0.8** | **68.7 ± 0.4** | **67.9 ± 1.2** | **30** |
| **34.2 ± 1.0** | **61.2 ± 0.7** | **68.1 ± 1.2** | **67.6 ± 1.5** | **60** |
| **33.7 ± 0.5** | **65.0 ± 1.4** | **68.4 ± 0.9** | **68.5 ± 0.5** | **300** |

| **POI attachment** | | | | |
|---|---|---|---|---|
| **Molar ratio DNA origami : bt-H57-scFv** | | | | |
| **1 : 1** | **1 : 5** | **1 : 10** | **1 : 100** | **Incubation time [min]** |
| **34.7 ± 2.2** | **59.8 ± 0.9** | **62.0 ± 0.4** | **66.9 ± 1.5** | **10** |
| **32.0 ± 1.2** | **59.1 ± 1.5** | **62.2 ± 1.3** | **67.5 ± 0.5** | **30** |
| **31.1 ± 1.4** | **60.1 ± 0.8** | **67.4 ± 1.0** | **68.1 ± 0.7** | **60** |
| **34.6 ± 1.0** | **66.6 ± 0.9** | **67.3 ± 1.0** | **67.5 ± 0.5** | **300** |

**Table S4. Optimization of functionalization conditions for construct H57-dSAv-NL.** For each step, the yield of functionalized DNA origami construct was determined for different conditions (molar ratios, incubation times) via two-color colocalization microscopy as sketched in SI Fig. S4. Cumulative yields are shown. Optimal conditions for each step are marked in red. Data are the mean (± sem) of at least two independent experiments.

| Handle incorporation | | | | | |
|---|---|---|---|---|---|
| Molar ratio DNA origami : biotinylated handle oligo | | | | | |
| 1 : 1 | 1 : 2 | 1 : 5 | 1 : 10 | 1 : 30 | 1 : 100 |
| 21.2 ± 1.2 | 53.8 ± 7.0 | 59.0 ± 7.0 | 84.0 ± 2.7 | 79.9 ± 6.2 | 79.4 ± 0.5 |

| SAv attachment | | | | |
|---|---|---|---|---|
| Molar ratio DNA origami : dSAv | | | | |
| 1 : 1 | 1 : 5 | 1 : 10 | 1 : 100 | Incubation time [min] |
| 43.2 ± 2.7 | 44.6 ± 6.6 | 64.7 ± 2.8 | 66.5 ± 3.8 | 10 |
| 46.8 ± 4.6 | 55.4 ± 8.6 | 71.7 ± 3.3 | 67.8 ± 2.6 | 30 |
| 44.2 ± 1.9 | 67.4 ± 3.6 | 72.0 ± 2.6 | 68.0 ± 2.5 | 60 |
| 46.1 ± 8.2 | 66.3 ± 3.0 | 69.1 ± 4.2 | 70.1 ± 3.1 | 300 |

| POI attachment | | | | |
|---|---|---|---|---|
| Molar ratio DNA origami : bt-H57-scF$_V$ | | | | |
| 1 : 1 | 1 : 5 | 1 : 10 | 1 : 100 | Incubation time [min] |
| 58.4 ± 4.8 | 63.5 ± 4.9 | 66.2 ± 4.5 | 68.5 ± 4.5 | 10 |
| 59.5 ± 5.7 | 66.1 ± 3.2 | 67.0 ± 5.0 | 69.6 ± 3.7 | 30 |
| 64.0 ± 2.4 | 66.4 ± 4.0 | 71.2 ± 2.6 | 69.8 ± 3.2 | 60 |
| 64.3 ± 5.3 | 67.4 ± 4.5 | 72.4 ± 0.3 | 73.2 ± 2.5 | 300 |

**Table S5. Optimization of functionalization conditions for construct H57-DNA.** For each step, the yield of functionalized DNA origami construct was determined for different conditions (molar ratios, incubation times) via two-color colocalization microscopy as sketched in SI Fig. S5. Cumulative yields are shown. Optimal conditions for each step are marked in red. Data are the mean (± sem) of at least two independent experiments.

| Handle incorporation | | | | | | |
|---|---|---|---|---|---|---|
| Molar ratio DNA origami : handle oligo | | | | | | |
| 1 : 1 | 1 : 2 | 1 : 5 | 1 : 10 | 1 : 30 | 1 : 50 | 1 : 100 |
| 19.1 ± 0.9 | 32.8 ± 0.8 | 46.6 ± 0.6 | 84.4 ± 2.1 | 83.4 ± 2.7 | 86.4 ± 0.3 | 86.4 ± 0.0 |

| POI attachment | | | | |
|---|---|---|---|---|
| Molar ratio DNA origami : H57-scFv-DNA | | | | |
| 1 : 1 | 1 : 5 | 1 : 10 | 1 : 100 | Incubation time [min] |
| 32.6 ± 2.6 | 62.8 ± 2.9 | 65.7 ± 1.9 | 67.1 ± 3.0 | 10 |
| 34.9 ± 4.3 | 63.7 ± 2.5 | 66.9 ± 2.4 | 67.6 ± 1.9 | 30 |
| 36.1 ± 3.3 | 65.4 ± 4.1 | 67.9 ± 2.4 | 67.6 ± 2.1 | 60 |
| 58.5 ± 1.7 | 66.3 ± 3.6 | 67.7 ± 1.8 | 67.8 ± 2.6 | 300 |

**Table S6. Determination of functionalization yields after each step for construct H57-PNA.** For each step, the yield of functionalized DNA origami construct was determined for different conditions (incubation times and molar ratios) via two-color colocalization microscopy as sketched in SI Fig. S6. Cumulative yields are shown. Optimal conditions for each step are marked in red. Data are the mean (± s.e.m.) of at least two independent experiments.

| **Handle incorporation** | | | | | | |
|---|---|---|---|---|---|---|
| **Molar ratio DNA origami : handle oligo** | | | | | | |
| **1 : 1** | **1 : 2** | **1 : 5** | **1 : 10** | **1 : 30** | **1 : 50** | **1 : 100** |
| **19.1 ± 0.9** | **32.8 ± 0.8** | **46.6 ± 0.6** | **84.4 ± 2.1** | **83.4 ± 2.7** | **86.4 ± 0.3** | **86.4 ± 0.0** |

| **POI attachment** | | | | |
|---|---|---|---|---|
| **Molar ratio DNA origami : H57-scFv-PNA** | | | | |
| **1 : 1** | **1 : 3** | **1 : 10** | **1 : 100** | **Incubation time [min]** |
| **24.7 ± 1.9** | **28.0 ± 1.5** | **47.9 ± 3.2** | **48.9 ± 3.2** | **10** |
| **50.2 ± 6.0** | **71.5 ± 2.5** | **71.7 ± 2.3** | **73.1 ± 2.4** | **30** |
| **52.0 ± 4.5** | **74.2 ± 2.2** | **72.9 ± 2.5** | **73.0 ± 3.0** | **60** |
| **51.4 ± 5.0** | **73.6 ± 2.0** | **72.9 ± 2.5** | **72.9 ± 2.7** | **300** |

**Table S7. Determination of functionalization yields after each step for construct H57-mSAv.** For each step, the yield of functionalized DNA origami construct was determined for different conditions (incubation times and molar ratios) via two-color colocalization microscopy as sketched in SI Fig. S7. Cumulative yields are shown. Optimal conditions for each step are marked in red. Data are the mean (± s.e.m.) of at least two independent experiments.

.

| **Handle incorporation** | | | | | | |
|---|---|---|---|---|---|---|
| **Molar ratio DNA origami : handle oligo** | | | | | | |
| **1 : 1** | **1 : 2** | **1 : 5** | **1 : 10** | **1 : 30** | **1 : 50** | **1 : 100** |
| **19.1 ± 0.9** | **32.8 ± 0.8** | **46.6 ± 0.6** | **84.4 ± 2.1** | **83.4 ± 2.7** | **86.4 ± 0.3** | **86.4 ± 0.0** |

| **SAv attachment** | | | |
|---|---|---|---|
| **Molar ratio DNA origami : mSAv-DNA** | | | |
| **1 : 3** | **1 : 10** | **1 : 100** | **Incubation time [min]** |
| **68.8 ± 2.7** | **74.0 ± 2.5** | **76.4 ± 3.1** | **10** |
| **75.4 ± 0.4** | **75.7 ± 1.5** | **78.1 ± 0.6** | **30** |
| **81.0 ± 2.4** | **78.5 ± 2.7** | **78.2 ± 5.8** | **60** |
| **78.0 ± 0.6** | **80.0 ± 1.9** | **81.1 ± 3.0** | **300** |

| **POI attachment** | | | | |
|---|---|---|---|---|
| **Molar ratio DNA origami : bt-H57-scF$_V$** | | | | |
| **1 : 1** | **1 : 5** | **1 : 10** | **1 : 100** | **Incubation time [min]** |
| **47.4 ± 3.0** | **50.0 ± 4.8** | **56.4 ± 6.3** | **62.1 ± 2.6** | **10** |
| **49.5 ± 1.5** | **58.3 ± 5.0** | **63.2 ± 2.4** | **65.5 ± 3.0** | **30** |
| **50.7 ± 1.6** | **63.1 ± 1.9** | **66.6 ± 3.2** | **66.3 ± 2.9** | **60** |
| **53.7 ± 1.7** | **65.5 ± 3.2** | **66.5 ± 3.1** | **67.6 ± 3.8** | **300** |

**Table S8. Determination of functionalization yields after each step for construct H57-tSAv.** For each step, the yield of functionalized DNA origami construct was determined for different conditions (incubation times and molar ratios) via two-color colocalization microscopy as sketched in SI Fig. S8. Cumulative yields are shown. Optimal conditions for each step are marked in red. Data are the mean (± s.e.m.) of at least two independent experiments.

**Handle incorporation**

**Molar ratio DNA origami : handle oligo**

| 1 : 1 | 1 : 2 | 1 : 5 | 1 : 10 | 1 : 30 | 1 : 50 | 1 : 100 |
|---|---|---|---|---|---|---|
| 19.1 ± 0.9 | 32.8 ± 0.8 | 46.6 ± 0.6 | 84.4 ± 2.1 | 83.4 ± 2.7 | 86.4 ± 0.3 | 86.4 ± 0.0 |

**bt-DNA hybridization**

**Molar ratio DNA origami : bt-DNA**

| 1 : 10 | 1 : 20 | 1 : 50 | 1 : 100 | 1 : 300 | 1 : 1000 |
|---|---|---|---|---|---|
| 26.9 ± 0.3 | 39.1 ± 2.0 | 86.4 ± 0.5 | 85.4 ± 0.3 | 85.4 ± 0.2 | 85.9 ± 0.0 |

**SAv attachment**

**Molar ratio DNA origami : tSAv**

| 1 : 1 | 1 : 5 | 1 : 10 | 1 : 100 | Incubation time [min] |
|---|---|---|---|---|
| 76.6 ± 4.8 | 77.9 ± 5.2 | 77.5 ± 5.7 | 79.0 ± 5.6 | 10 |
| 77.5 ± 3.6 | 79.7 ± 4.8 | 79.4 ± 3.2 | 79.4 ± 3.6 | 30 |
| 76.8 ± 4.3 | 78.3 ± 6.0 | 79.6 ± 4.0 | 79.3 ± 4.0 | 60 |
| 77.7 ± 4.9 | 78.8 ± 6.1 | 79.6 ± 4.0 | 82.1 ± 2.6 | 300 |

**POI attachment**

**Molar ratio DNA origami : bt-H57-scF$_V$**

| 1 : 10 | Incubation time [min] |
|---|---|
| 69.1 ± 2.3 | 10 |
| 70.5 ± 1.1 | 30 |
| 70.1 ± 1.3 | 60 |
| 68.1 ± 3.0 | 300 |

**Table S9. Determination of functionalization yields after each step for construct H57-tSAv-NL.** For each step, the yield of functionalized DNA origami construct was determined for different conditions (incubation times and molar ratios) via two-color colocalization microscopy as sketched in SI Fig. S9. Cumulative yields are shown. Optimal conditions for each step are marked in red. Data are the mean (± s.e.m.) of at least two independent experiments.

| Handle incorporation | | | | | |
|---|---|---|---|---|---|
| Molar ratio DNA origami : biotinylated handle oligo | | | | | |
| 1 : 1 | 1 : 2 | 1 : 5 | 1 : 10 | 1 : 30 | 1 : 100 |
| 21.2 ± 1.2 | 53.8 ± 7.0 | 59.0 ± 7.0 | 84.0 ± 2.7 | 79.9 ± 6.2 | 79.4 ± 0.5 |

| SAv attachment | | | | |
|---|---|---|---|---|
| Molar ratio DNA origami : tSAv | | | | |
| 1 : 1 | 1 : 5 | 1 : 10 | 1 : 100 | Incubation time [min] |
| 65.2 ± 6.5 | 78.8 ± 0.8 | 79.5 ± 1.4 | 81.2 ± 1.7 | 10 |
| 67.9 ± 10.5 | 79.5 ± 1.5 | 83.8 ± 1.3 | 81.3 ± 1.7 | 30 |
| 78.0 ± 2.4 | 80.6 ± 2.3 | 82.7 ± 0.3 | 81.5 ± 1.8 | 60 |
| 81.0 ± 1.4 | 81.6 ± 1.1 | 82.1 ± 2.4 | 83.9 ± 0.2 | 300 |

| POI attachment | |
|---|---|
| Molar ratio DNA origami : bt-H57-scF$_V$ | |
| 1 : 10 | Incubation time [min] |
| 71.0 ± 1.3 | 10 |
| 71.2 ± 1.2 | 30 |
| 72.1 ± 1.9 | 60 |
| 71.6 ± 1.8 | 300 |

**Table S10. Functionalization efficiencies for the different constructs.** The degree of functionalization (%, ± s.e.m.) (with one or more ligands) was determined via two-color colocalization microscopy (see Figures SI 3-9). Yields for individual steps *(A)* and cumulative yields *(B)* are shown.

| *A* | H57-dSAv | H57-dSAv-NL | H57-DNA | H57-PNA | H57-mSAv | H57-tSAv | H57-tSAv-NL |
|---|---|---|---|---|---|---|---|
| **handle incorporation** | 84.0 ± 2.7 | 84.0 ± 2.7 | 84.4 ± 2.1 | 84.4 ± 2.1 | 84.4 ± 2.1 | 84.4 ± 2.1 | 84.0 ± 2.7 |
| **bt-DNA hybridization** | - | - | - | - | - | 102.4 ± 3.1 | - |
| **SAv attachment** | 85.7 ± 3.0 | 85.7 ± 3.0 | - | - | 96.0 ± 5.2* | 91.9 ± 6.5 | 99.8 ± 4.8 |
| **H57-scFv attachment** | 98.9 ± 3.9 | 98.9 ± 3.9 | 80.5 ± 2.8* | 87.9 ± 2.5* | 82.2 ± 3.9 | 88.3 ± 7.4 | 86.0 ± 2.2 |
| ***B*** | **H57-dSAv** | **H57-dSAv-NL** | **H57-DNA** | **H57-PNA** | **H57-mSAv** | **H57-tSAv** | **H57-tSAv-NL** |
| **handle incorporation** | 84.4 ± 2.1 | 84.0 ± 2.7 | 84.4 ± 2.1 | 84.4 ± 2.1 | 84.4 ± 2.1 | 84.4 ± 2.1 | 84.0 ± 2.7 |
| **bt-DNA hybridization** | 86.4 ± 0.5 | - | - | - | - | 86.4 ± 0.5 | - |
| **SAv attachment** | 68.7 ± 0.1 | 72.0 ± 0.2 | - | - | 81.0 ± 2.4* | 79.4 ± 3.2 | 83.8 ± 1.3 |
| **H57-scFv attachment** | 67.4 ± 0.7 | 71.2 ± 2.6 | 67.9 ± 0.7* | 74.2 ± 0.3* | 66.6 ± 1.2 | 70.1 ± 0.9 | 72.1 ± 0.7 |

*indicates attachment via hybridization

**Table S11. Number of H57-scF$_V$ molecules per DNA origami construct.** The number of H57-scF$_V$ molecules per functionalized DNA origami construct was determined by comparing the signal brightness of the construct to the brightness of a single AF555-labeled H57-scF$_V$ as described in the methods section. Data are the mean of at least two independent experiments (± s.e.m.).

| Construct | 1 H57-scF$_V$ | 2 H57-scF$_V$ | 3 H57-scF$_V$ | signals (n) |
|---|---|---|---|---|
| **H57-dSAv** | 98.6 ± 0.8 | 0.5 ± 0.3 | 0.6 ± 0.5 | 1,352 ± 147 |
| **H57-dSAv-NL** | 99.9 ± 0.3 | 0.1 ± 0.1 | 0.6 ± 0.2 | 1,369 ± 107 |
| **H57-DNA** | 97.5 ± 1.1 | 2.2 ± 1.1 | 0.3 ± 0.2 | 1,273 ± 69 |
| **H57-PNA** | 99.5 ± 0.2 | 0.0 ± 0.0 | 0.5 ± 0.2 | 2,276 ± 484 |
| **H57-mSAv** | 98.6 ± 0.8 | 1.1 ± 0.7 | 0.3 ± 0.1 | 1,359 ± 63 |
| **H57-tSAv** | 30.6 ± 1.8 | 52.8 ± 3.2 | 16.7 ± 5.1 | 5,984 ± 1,627 |
| **H57-tSAv-NL** | 48.1 ± 3.2 | 47.1 ± 0.7 | 4.8 ± 3.9 | 3,096 ± 565 |

**Table S12. Fitting parameters of fits to dose-response curves.** Dose-response curves of T-cell activation by via the different ligand-decorated DNA origami constructs were fitted with Eq. 19 to extract the activation threshold $T_A$, the maximum response $A_{max}$ and the Hill coefficient n. The 95% confidence intervals are indicated. The mean number of cells per region (± s.e.m.) and the number of animals used to generate dose-response curves are shown.

| Construct | n | $n_{low}$ | $n_{high}$ | $A_{max}$ (%) | $A_{max, low}$ | $A_{max, high}$ | $T_A$ ($\mu m^{-2}$) | $T_{A, low}$ | $T_{A, high}$ | # cells | # animals |
|---|---|---|---|---|---|---|---|---|---|---|---|
| **H57-dSAv** | 2.44 | 1.66 | 3.23 | 91.81 | 84.50 | 99.13 | 3.91 | 3.29 | 4.64 | 163 ± 35 | 3 |
| **H57-dSAv-NL** | 3.06 | 1.83 | 4.29 | 100.14 | 91.55 | 108.73 | 3.27 | 2.78 | 3.85 | 199 ± 55 | 2 |
| **H57-DNA** | 4.38 | 2.49 | 6.26 | 91.01 | 81.41 | 100.62 | 9.68 | 8.66 | 10.83 | 176 ± 68 | 2 |
| **H57-PNA** | 1.73 | 0.94 | 2.52 | 103.83 | 94.52 | 113.15 | 2.75 | 2.20 | 3.43 | 206 ± 65 | 2 |
| **H57-mSAv** | 3.35 | 0.99 | 5.71 | 92.55 | 84.01 | 101.09 | 3.15 | 2.58 | 3.86 | 203 ± 53 | 2 |
| **H57-tSAv** | 1.23 | 0.33 | 2.12 | 105.74 | 89.16 | 122.31 | 0.81 | 0.45 | 1.49 | 289 ± 81 | 2 |
| **H57-tSAv-NL** | 1.42 | 0.72 | 2.12 | 101.65 | 90.28 | 113.02 | 0.83 | 0.45 | 1.54 | 214 ± 62 | 2 |

**Table S13. List of staple strands**

| Designation | Sequence |
| --- | --- |
| 10[63]-BLK | CAGCTTTCGGGCGCATCGTAACCGATCGGCCT |
| 11[112]-BLK | AGCGCGAACAATCATAAGGGAACCCGGTGTAC |
| 12[95]-BLK | ACACTCATAGGGGACGACGACAGTTGCATCTG |
| 13[80]-BLK | CGCCAGCTAAAGACAGCATCGGAAGTCACCCT |
| 14[63]-BLK | GCGATTAAGTACCGAGCTCGAATTAATTGTTA |
| 15[112]-BLK | TTTCTTAACCGCTTTTGCGGGATCCGAGGGTA |
| 16[95]-BLK | GTATCGGTGCTGTTTCCTGTGTGACGTAATCA |
| 19[112]-BLK | TAGCAAGCCGATCTAAAGTTTTGTGTATGGGA |
| 2[63]-BLK | TATTTTCATTAAGCAATAAAGCCTACATTATG |
| 5[80]-BLK | CGGAGACATTCATCAGTTGAGATTATTACAGG |
| 6[63]-BLK | TCAACCGTATCGATGAACGGTAATGGTTGATA |
| 7[112]-BLK | TAATTTCACTAACGGAACAACATTTAGGAATA |
| 8[95]-BLK | AGAACGAGCAATCATATGTACCCCCGTAAAAC |
| 9[80]-BLK | CAAAAATAAAAGAGGACAGATGAAGAACTGAC |
| 1[32]-BLK | TGATTCCCAAAAGGTGGCATCAATAATCATAC |
| 17[32]-BLK | CATTAATTGGGCGCCAGGGTGGTTATTGCCCT |
| 20[143]-BLK | AGGAGGTTTACCGTAACACTGAGTCATTCCAC |
| 4[143]-BLK | TACCAGACCGGAATCGTCATAAATGTTCAGAA |
| 0[143]-BLK | ACAGGTCAGGATTAGAGAGTACCTGAAGCCCG |
| 0[47]-BLK | GCTCAACATGTTTTAAATATGCAAATAACAGT |
| 0[63]-BLK | TGAATATAAGATTTAGTTTGACCATTTAGCTA |
| 0[95]-BLK | TCATTTTTGCGGATGGCTTAGAGCTTAATTGC |
| 1[112]-BLK | ATTAAGAGTTAATTGCTCCTTTTGATAAGAGG |
| 1[128]-BLK | AAAGACTTTGACCATAAATCAAAAATAGCGTC |
| 1[32]-BLK | TGATTCCCAAAAGGTGGCATCAATAATCATAC |
| 1[80]-BLK | ATTTCGCAAGCAAAGCGGATTGCAGACTATTA |
| 10[143]-BLK | TAGCCGGACCTTCATCAAGAGTAATCAACGTA |
| 10[95]-BLK | CAACTTTGATTCGCGTCTGGCCTTAACGCCAT |
| 11[32]-BLK | GGGATAGGTTTCCGGCACCGCTTCCATTCAGG |
| 11[80]-BLK | CCAGTTTGCTTTGACCCCCAGCGAACACTAAA |
| 12[143]-BLK | CTACGAAGATTTGTATCATCGCCTATGTTACT |
| 12[47]-BLK | CAGCCAGCTCACGTTGGTGTAGATATCAACAT |
| 12[63]-BLK | CAGGAAGATCGGTGCGGGCCTCTTGCTGCAAG |
| 13[112]-BLK | GCAACGGCAAAGAGGCAAAAGAATTTATACCA |
| 13[128]-BLK | CTTTGAGGTGCAGGGAGTTAAAGGACAGCTTG |
| 13[32]-BLK | CTGCGCAAGTTTTCCCAGTCACGAATGCCTGC |
| 14[143]-BLK | CTGAGGCTACTAAAGACTTTTTCATAATGCCA |
| 14[95]-BLK | CAGCAGCGGGCGAAAGGGGGATGTCGCTATTA |
| 15[32]-BLK | AGGTCGACACGAGCCGGAAGCATACTAACTCA |
| 15[80]-BLK | TGGTCATATTATCAGCTTGCTTTCCTTTAATT |
| 16[143]-BLK | TCACGTTGAGTTGCGCCGACAATGTTCGGTCG |
| 16[47]-BLK | CACAACATTCTAGAGGATCCCCGGGTTGGGTA |
| 16[63]-BLK | TCCGCTCACCGCTTTCCAGTCGGGGCCAACGC |
| 17[112]-BLK | TTTTGCTAAGGCTCCAAAAGGAGCGAGGTGAA |
| 17[128]-BLK | TCAACAGTCTCATAGTTAGCGTAACCAATAGG |
| 17[32]-BLK | CATTAATTGGGCGCCAGGGTGGTTATTGCCCT |
| 17[80]-BLK | CGTGCCAGAGTAAATGAATTTTCTCGTCTTTC |

| | |
|---|---|
| 18[143]-BLK | AGACAGCCTTCAGCGGAGTGAGAATAATTTTT |
| 18[63]-BLK | GCGGGGAGGCAGCAAGCGGTCCACTGATGGTG |
| 18[95]-BLK | CAGACGTTCTGCATTAATGAATCGAAACCTGT |
| 19[32]-BLK | TCACCGCCATAAATCAAAAGAATAGGAACAAG |
| 19[80]-BLK | GCCCCAGCAGCCACCACCCTCATTGAACCGCC |
| 2[143]-BLK | AACGAGAACAAATATCGCGTTTTAAAACTCCA |
| 2[95]-BLK | TAGTCAGAAATGGTCAATAACCTGTTAGATAC |
| 20[143]-BLK | AGGAGGTTTACCGTAACACTGAGTCATTCCAC |
| 20[47]-BLK | AATCCCTTTGGCCCTGAGAGAGTTAGGCGGTT |
| 20[63]-BLK | GTTCCGAATCCAACGTCAAAGGGCGAAAAACC |
| 20[95]-BLK | ACCCTCAGAGGCGAAAATCCTGTTGCTGGTTT |
| 21[112]-BLK | TTTTGCTCAGAACCGCCACCCTCATTCAGGGA |
| 21[128]-BLK | GCGGATAAGTGCCGTCGAGAGGGTCCGTACTC |
| 21[32]-BLK | AGTCCACTATTAAAGAACGTGGACATCGGCAA |
| 21[80]-BLK | GTCTATCAAGAGAAGGATTAGGATTAGCGGGG |
| 3[112]-BLK | TAGACTGGATCAGGTCTTTACCCTTCAAAAAG |
| 3[32]-BLK | AGGCAAGGAGCCTTTATTTCAACGTTTAAATG |
| 3[80]-BLK | AAAGCTAACAGAGGGGGTAATAGTGCAAAAGA |
| 4[143]-BLK | TACCAGACCGGAATCGTCATAAATGTTCAGAA |
| 4[47]-BLK | GCGGGAGACAAAGAATTAGCAAAATTTGGGGC |
| 4[63]-BLK | ACCCTGTAAAGATTCAAAAGGGTGTATGATAT |
| 4[95]-BLK | AGTTTTGCATCGGTTGTACCAAAACAGAGCAT |
| 5[112]-BLK | CCACATTCATAGCGAGAGGCTTTTAAAATGTT |
| 5[128]-BLK | CAGATACAACGTTAATAAAACGAAACTTTAAT |
| 5[32]-BLK | CAATGCCTATGCCGGAGAGGGTAGTCATTGCC |
| 6[143-BLK] | AAAAATCTTAACGCCAAAAGGAATCCCTCGTT |
| 6[95]-BLK | TAGAAAGAGTCAAATCACCATCAAAGAAAGGC |
| 7[32]-BLK | TGAGAGTCAGATTGTATAAGCAAAAATTCGCA |
| 7[80]-BLK | TAGCATGTTAGTAAATTGGGCTTGGAAACACC |
| 8[143-BLK] | ACAAAGCTATTACCTTATGCGATTTTGGGAAG |
| 8[47]-BLK | AAACAGGATGGAGCAAACAAGAGATCTAGCTG |
| 8[63]-BLK | ATCAGAAATTTTTTAACCAATAGGCCTGTAGC |
| 9[112-BLK] | AGACCAGGAGGCTTGCCCTGACGAAGATGGTT |
| 9[128-BLK] | CTGGCTGAACGAGGCGCAGACGGTACAAAGTA |
| 9[32]-BLK | TTAAATTTAGCGAGTAACAACCCGACCGTAAT |

**Table S14. List of elongated staple strands**

| Designation | Sequence | Docking Sequence |
|---|---|---|
| 9[80]-4T-V' | CAAAAATAAAAGAGGACAGATGAAGAACTGACTTTTGTGGAGTAGTGTCATGT | GTGGAGTAGTGTCATGT |
| Z'-4T-10[47] | AGAGTCCTAGCATATTTAGCCTTTTTAAATGTGTTGTTAAATCAGCTCAAGCCCCAA | AGAGTCCTAGCATATTTAGCC |
| Z'-4T-11[128] | AGAGTCCTAGCATATTTAGCCTTTTCAACGGAGGCACCAACCTAAAACGTACAGAGG | AGAGTCCTAGCATATTTAGCC |
| Z'-4T-14[47] | AGAGTCCTAGCATATTTAGCCTTTTACGCCAGGCTGTTGGGAAGGGCGATCGCACTC | AGAGTCCTAGCATATTTAGCC |
| Z'-4T-15[128] | AGAGTCCTAGCATATTTAGCCTTTTATACCGATAAAATCTCCAAAAAAAAACAACTT | AGAGTCCTAGCATATTTAGCC |
| Z'-4T-18[47] | AGAGTCCTAGCATATTTAGCCTTTTTGCGTATTGCGTTGCGCTCACTGCCAATTCCA | AGAGTCCTAGCATATTTAGCC |
| Z'-4T-19[128] | AGAGTCCTAGCATATTTAGCCTTTTAACCCATGTAGTACCGCCACCCTCAGTACCAG | AGAGTCCTAGCATATTTAGCC |
| Z'-4T-2[47] | AGAGTCCTAGCATATTTAGCCTTTTGCGAGCTGAATTCTGCGAACGAGTATGCTGTA | AGAGTCCTAGCATATTTAGCC |
| Z'-4T-3[128] | AGAGTCCTAGCATATTTAGCCTTTTCAATACTGGACGATAAAAACCAAAAACTAATG | AGAGTCCTAGCATATTTAGCC |
| Z'-4T-6[47] | AGAGTCCTAGCATATTTAGCCTTTTATAAATTAGAGTAATGTGTAGGTAATACTTTT | AGAGTCCTAGCATATTTAGCC |
| Z'-4T-7[128] | AGAGTCCTAGCATATTTAGCCTTTTCATTGTGAGCTCATTCAGTGAATACGCATAGG | AGAGTCCTAGCATATTTAGCC |

**Table S15. List of modified oligonucleotides**

| Designation | Composition | Sequence | Modification |
|---|---|---|---|
| bt-DNA | Biotin-TEG-V | ACATGACACTACTCCAC | 5'-Biotin-TEG |
| bt-DNA-635P | Biotin-TEG-V-4T-AS635P | ACATGACACTACTCCACTTTT | 5'-Biotin-TEG; 3'- AS635P |
| DNA-635P | 9[80]-4T-V'-AS635P | CAAAAATAAAAGAGGACAGATGAAGAACTGACTTTTGTGGAGTAGTGTCATGT | 3'- AS635P |
| AF647-DNA-bt | AF647-2T-9[80]-2T-TEG-Biotin | TTCAAAAATAAAAGAGGACAGATGAAGAACTGACTT | 5'- AF647 ; 3'- Biotin-TEG |
| mSAv-DNA | mSAv-PEG5-V | TTTTACATGACACTACTCCAC | 5'- mSA-TEG |
| H57-DNA | H57-PEG5-4T-V | TTTTACATGACACTACTCCAC | 5'- H57-scF$_V$-PEG5 |
| H57-PNA | H57-2T-V (PNA) | O - TTACATGACACTACTCCAC | 5'- H57-scF$_V$-O-linker |
|  | Z-TEG-Cholesterol | GGCTAAATATGCTAGGACTCT | 3'-Cholesterol-TEG |
